# Unravelling the memory of the extracellular matrix using MASH-derived decellularized scaffolds

**DOI:** 10.64898/2026.03.17.712486

**Authors:** Gabriel Reis Pinto, Luana Diniz Guerra Braz, Yasmin Pestana, Alexandre Cerqueira da Silva Filho, Maria Isabel Moraes do Amaral Candido Gomes, Julia Helena Oliveira de Barros, Thamires Siqueira de Oliveira, Isadora Z.L.F. Feng, Barbara Fidelix Santana, Hernandes F Carvalho, Cherley Borba Oliveira de Andrade, Lucas Pires Guarnier, Érica Almeida Amorim, Cibele Ferreira Pimentel, Alfredo M. Goes, Maria de Fátima Leite, Robson A.S. Santos, Marina Amaral Alves, Regina Coeli dos Santos Goldenberg, Marlon Lemos Dias

## Abstract

The use of decellularized diseased livers in regenerative medicine is a promising approach for eliminating organ shortages. Bioengineering studies have shown that ECM can impact cell physiology, inducing cell activation, function, and ECM deposition, which suggests that the ECM has a “memory” that is involved in the outcome after recellularization. However, the effect of diseased ECM memory on new cells in vitro and in vivo has not been thoroughly investigated. Since it has been increasingly recognized that liver ECM changes due to different factors, it is comprehensively that diseased ECM obtained from discarded organs will ensure a distinct environment and impact cell survival and physiology. Thus, we aimed at investigating the impact of the memory of diseased ECM obtained from metabolic dysfunction-associated steatohepatitis (MASH)-derived organs on steatohepatitis establishment. To address this aim, we explored decellularized ECM obtained from rats and humans with MASH in different contexts. First, MASH ECM was characterized and then submitted to transplantation to investigate whether a MASH-derived ECM could be used as a scaffold for transplantation and to promote steatohepatitis features in control animals. Histological analysis revealed that the MASH-ECM was completely recellularized after transplantation in both control and MASH recipient rats. However, steatosis and fibrosis were observed in MASH ECM after transplantation in both groups. Molecular analysis showed that MASH ECM stimulates de novo lipogenesis and fibrosis 30 days after transplantation. Untargeted metabolomic analysis revealed that cells grown on MASH ECM had a similar metabolic profile, even when transplanted into healthy or MASH recipient rats. In addition, we observed that MASH ECM promoted impaired lipid oxidation and mitochondrial dysfunction when transplanted into healthy recipients. Altered lipid turnover and inflammatory signaling were observed in MASH ECM transplanted in MASH recipients. In vitro analysis revealed that MASH ECM induced lipid accumulation in HepG2 cells after 10 days of culture. Calcium signalling experiments obtained from HepG2 cells cultured in MASH ECM showed a lower response to ATP, a reduced calcium signalling amplitude, and a distinct response profile than that observed in healthy ECM. On the other hand, a diseased human-derived ECM could still provide an environment that allows cell development. Taken together, our data showed that MASH ECM impacts cell metabolism, promoting steatohepatitis maintenance. In conclusion, our data confirm that diseased ECM memory can impact cell physiology contributing to disease progression.

## Introduction

Liver diseases account for 4% of all deaths annually, comprising over two million deaths worldwide [1]. Liver transplantation (LT) represents the most effective life-saving treatment for end-stage liver disease (ESLD), and the increasing burden of ESLD heightens the importance of this treatment for chronic liver disease (CLD) patients management [2]. LT history began in 1967, when the first successful deceased donor LT was performed by Thomas Starzl and colleagues, and since then, the success rates of this procedure have steadily risen over the decades through innovations in organ preservation strategies, surgical techniques, perioperative care, and immunosuppression [3, 4]. However, the discrepancy between the increased demand for organs and the available supply from deceased donors created an organ shortage scenario reflected in large transplantation waiting lists, in which insufficient functioning results in high mortality rates of patients depending on transplants [3, 4, 5]. To address this disparity, various alternatives are being scientifically explored, including the development of acellular liver scaffolds (ALS) based on decellularized ECMs derived from decellularization techniques [7].

The in vivo ALS exploration burden began in 2015, when the first successful orthotopic transplantation of ALS into mice was reported, confirming that descellularized matrices can be transplanted into healthy recipient animals [8]. This was followed by the successful transplantation of ALS in rats after partial hepatectomy, highlighting the availability of in vivo recellularization of decellularized grafts [9]. In addition to the observation of the interaction between cells and the ECM of healthy scaffolds, In vitro experiments reported by Miyauchi and colleagues showed that the decellularized ECM obtained from cirrhotic rat livers promotes the acceleration of the progression of hepatocellular carcinoma cells to a more aggressive and proliferative profile [10]. Similar results were observed by Mazza and colleagues when human cirrhotic scaffolds were recellularized with HepG2 cells [11]. In addition to the liver, the influence of a fibrotic ECM on cells has also been observed in scaffolds originating from other organs. Booth and colleagues also observed that fibroblasts were activated and differentiated, acquiring a myofibroblast phenotype when cultured in human fibrotic lung scaffolds [12]. In this context, the combination of biochemical and structural components of the ECM can alter the phenotype and genotype of the cell. In 2023, data from our group in rat models showed that decellularized liver ECM derived from healthy animals can be successfully transplanted into cirrhotic recipient rats, enabling in vivo regeneration through stimulating recruitment and proliferation of endogenous cells [7]. Interestingly, it was found that the transplanted healthy liver ECM was not affected by the diseased liver and created an environment that stimulated healthy cell growth, regeneration, and repair [7]. Together, all these data pointed out the peculiar characteristics of ECM suggested by us as “ECM memory”. The extracellular matrix encodes biochemical and structural signals, in addition to matrix-bound molecules, which might include growth factors, that shape cellular genotype, phenotype, and physiology. These ECM-mediated cues or “zip codes” direct the behavior of resident and interacting cells, such that homeostatic matrix properties promote normal tissue development and function [13, 14], whereas altered or diseased matrix states drive maladaptive cellular responses and diseased tissue formation [10, 12, 15].

Taking into account the clinical reality, the possible source of organs for ALS generation would be discarded marginal livers, that represents organs that do not have any potential direct application for transplantation [7, 8, 16]. Epidemiological data have shown that the global prevalence of metabolic dysfunction-associated steatotic liver disease (MASLD) in the period between 1990-2019 was about 30.1%, representing an increase of 50.4% over the past 3 decades [17, 18]. In the period comprising 2002-2019, metabolic dysfunction-associated steatohepatitis (MASH) was the most non-hepatocellular carcinoma (HCC) cause of CLD among patients in the LT waiting list registered in the Scientific Registry of Transplant Recipients (SRTR) system [17, 19]. Considering the important contribution of MASLD and MASH complications for the rising in LT indication rates, it is possible to hypothesize that steatotic marginal livers derived from ESLD patients discarded after LT could be a relevant organ source for ALS manufacturing, with the finality of further being utilized for the development of artificial bioengineered organs, that could enable a solution for the liver supply shortage issue [16, 17]. Nevertheless, bioengineering approaches for the development of functional ALS derived from diseased organs are still under investigation, and the impact of the transplantation of diseased-derived ALS in diseased recipients is unknown. Thus, to continue the progress on the development of this promising technology for LT-indicated patients’ treatment it is necessary to investigate whether a diseased ECM can be used as an ALS in transplantation. In addition, in this study we also sought to expand the understanding of ECM memory through a new set of questions: a) whether a MASH ECM can be transplanted into a diseased recipient; b) whether a MASH ECM can be recellularized when transplanted into a diseased recipient; c) whether the diseased ECM impact in recipient liver metabolism, functional and injury marks; d) whether a MASH ECM can retain memory and impact on in vivo recellularization and promote steatohepatitis; e) whether a MASH ECM can impact on healthy cells phenotype inducing steatohepatitis-like features.

## Materials and Methods

### 2.1 Experimental design

Liver steatohepatitis was induced in female Wistar rats through high-fat diet administration in association with sucrose and intraperitoneal injections of carbon tetrachloride (CCl_4_) (see section 2.3 below). Steatohepatitis-induced animals were divided into two cohorts, donor and recipient rats (Figure 1). Donor animals underwent a total hepatectomy procedure. Excised livers were decellularized to obtain ECM MASH. Recipient animals were submitted to a partial hepatectomy of the median lobe (10%) and a MASH ECM fragment was transplanted into the excised site. Healthy animals were also submitted to MASH ECM orthotopic transplantation. After 30 days, recipient animals were euthanized. Transplanted liver and recellularized MASH ECM samples were submitted to multiple analyses, including histology, RT-qPCR, metabolomics, *in vitro* and *ex vivo* analysis. Human MASH/HCC ECM were also used in vitro to explore the memory of ECM.

**Figure 1.**
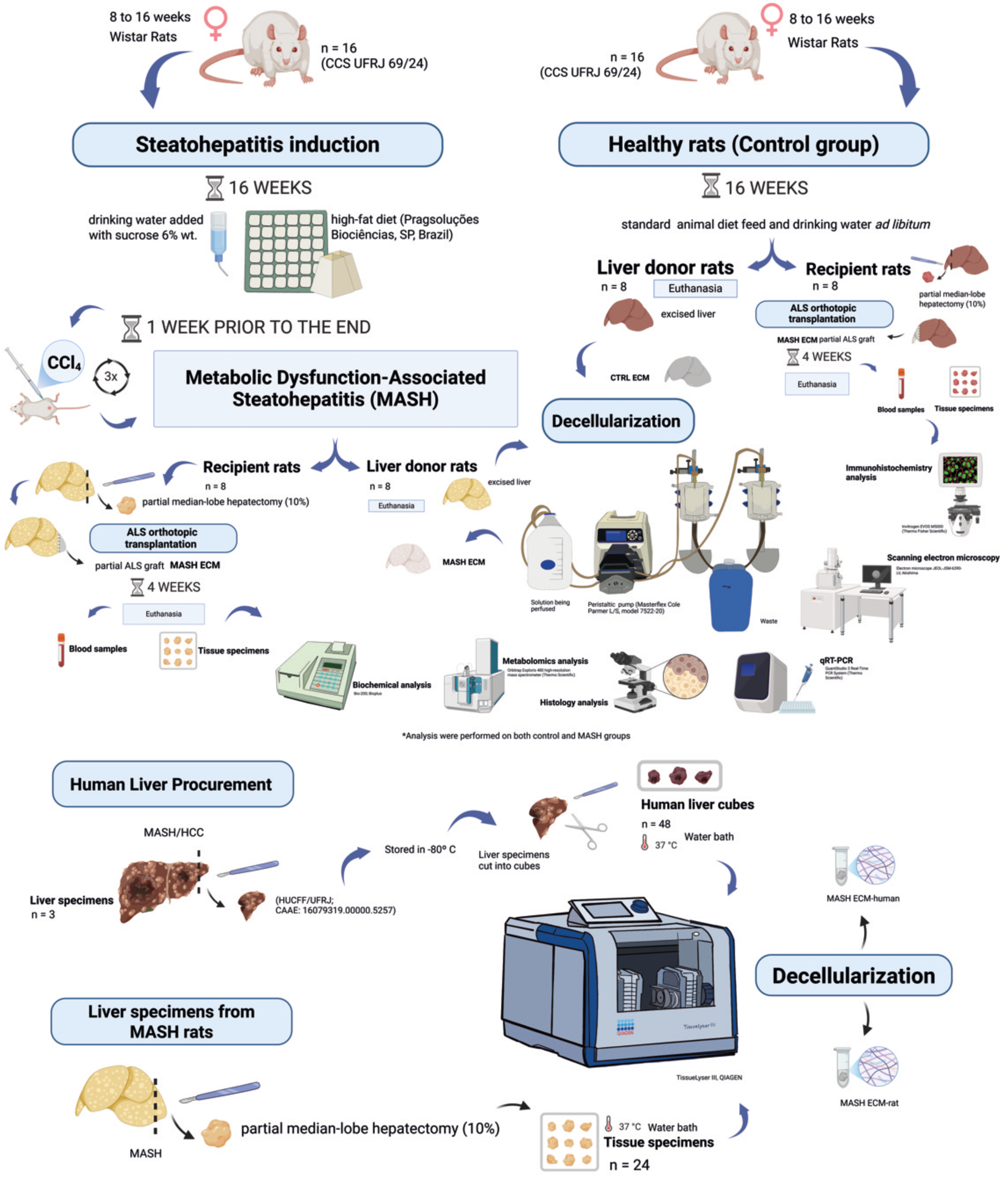
Experimental design. Liver steatohepatitis was induced in female Wistar rats through high-fat diet administration in association with intraperitoneal injections of carbon tetrachloride (CCl_4_). The steatohepatitis-induced animals were divided into two cohorts: donor and recipient rats. The donor animals underwent total hepatectomy. The excised livers were decellularized to obtain ECM MASH. Recipient animals underwent partial hepatectomy of the median lobe (10%), and a MASH ECM fragment was transplanted into the excised site. Healthy animals were also subjected to MASH-ECM orthotopic transplantation. After 30 days, the recipient animals were euthanized. Transplanted liver and recellularized MASH ECM samples were subjected to multiple analyses, including histology, qRT-PCR, and *in vitro* and *ex vivo* analyses. Human MASH/HCC ECM was also used in vitro to explore ECM memory. Liver specimens derived from MASH rats and human MASH/HCC were decellularized based on the procedure described by Thanapirom et al. (2021).

### 2.2 Animals

All animal procedures carried out in this study were approved and followed the animal care guidelines of the Animal Use Ethics Committee of the Health Science Center of the Federal University of Rio de Janeiro, Brazil (CEUA CCS UFRJ 69/24) (Figure 1). Female Wistar rats, ranging in age from 8 to 16 weeks, were randomly assigned into the following groups: MASH liver donor rats (n=8), healthy liver donor rats (n=8), ALS recipient rats (n=16) and healthy recipient rats (n=8). Animals were kept on a 12 h light/dark cycle, 25 ◦C mean ambient temperature, and 55 +/-5% humidity. The rats were fed with a high-fat diet, specific for steatohepatitis induction, and water *ad libitum.* Their body weight was monitored weekly. Anesthesia was induced by inhalation of 3-4% isoflurane (Isoforine®, Cristália, São Paulo, Brazil) and maintained by inhalation of 1-2% isoflurane, associated with an oxygen dose of 0.3-0.5 L/min.

### 2.3 Steatohepatitis induction

Steatohepatitis was induced by a high-fat diet containing 45% kcal from palm oil derived fat, 20% kcal from fructose, and 2% cholesterol (Pragsoluções Biociências, São Paulo, Brazil) to female Wistar rats aged from 8 to 16 weeks for 4 months. In addition to the diet, the animals received 6% sucrose in their drinking water *ad libitum* throughout the entire protocol (Figure 1). One week prior to the ending, animals received intraperitoneal injections of carbon tetrachloride (CCl₄; 1 mL/kg in olive oil, 1:1) on alternate days for a total of three injections.

### 2.4. Liver Procurement

#### ·2.4.1 Rat Liver Procurement

MASH and healthy donor animals (n=8 each) received an intraperitoneal injection of heparin (100 IU; Hemofol®, Cristália, São Paulo, Brazil) prior to the surgical procedure. Euthanasia was then performed by inhalational anesthetic overdose using isoflurane (2% isoflurane, associated with an oxygen dose of 1-2 L/min). A transverse abdominal incision followed by laparotomy was executed and the portal vein (PV) was identified, isolated, and cannulated with a 24-gauge catheter (Angiocatch®, BD, São Paulo, Brazil). The liver was subsequently excised and transferred to a sterile 60-mm Petri dish for subsequent decellularization process (Figure 1).

#### ·2.4.2 Human Liver Procurement

Liver specimens were obtained from patients with hepatocellular carcinoma arising from the setting of metabolic dysfunction–associated steatohepatitis (MASH). Those samples were from patients who underwent a curative intent surgical resection or from patients who underwent liver transplantation. The use of liver specimens was approved by the medical ethical council of the Clementino Fraga Filho University Hospital (HUCFF/UFRJ; CAAE: 16079319.00000.5257). After tissue collection, liver specimens were washed with saline solution 3 times for 5 min each and then stored in -80C until decellularization steps.

### 2.5 Decellularization

#### ·2.5.1 MASH rat livers decellularization

MASH-rat livers (n=8) were decellularized using a peristaltic pump (Masterflex Cole Parmer L/S, model 7522-20) at a flow rate of 7 mL/min. Livers were first subjected to continuous perfusion with distilled water for 2 h, followed by perfusion of 1% (v/v) Triton X-100 (Sigma-Aldrich, Saint Louis, MO, USA) through the PV for 2 h. Subsequently, a 1% (w/v) sodium dodecyl sulfate (SDS; Synth, São Paulo, Brazil) solution was perfused for 48 h. At the end of the process, ECM MASH-rat livers were perfused with distilled water overnight, followed by sterilization: 0.1% (v/v) sterile peracetic acid (PAA; SigmaAldrich 433241) at 3 mL/min for 1 h and by two washing cycles with 1% (v/v) sterile phosphate-buffered saline (PBS; Nova Biotecnologia 13-30262-05; São Paulo, Brazil) and stored in 0.1% (v/v) sterile PAA (SigmaAldrich 433241) at 4 °C until transplantation.

#### ·2.5.2 Human liver decellularization

The decellularization protocol was based on the procedure described by Thanapirom et al. (2021) [20]. First, liver specimens were cut into cubes (n=48) and stored individually at -80 C in 2 mL microtubes. They were then subjected to a water bath at 37 °C for 45 min. The microtubes were then filled with approximately 1.5 mL of 1X phosphate-buffered saline (PBS) and subjected to a water bath again for 15 min at 37 °C. After complete thawing, human liver cubes were subjected to the physical and chemical decellularization process using Tissue Lyser III (QIAGEN), which consists of short agitation cycles with a frequency from 15 to 20 Hz. During cycles, the microtubes were filled with 1.5 mL of ultrapure water, alternating with 1.5 mL of a specific RM (reagent mix) decellularization solution that consists of 4.3% (w/v) sodium chloride, 0.5% (w/v) sodium lauryl sulfate, 0.003% (v/v) Triton X-100, 3% (w/v) sodium deoxycholate and trypsin.EDTA 0.0025% (v/v). To verify the efficiency of the decellularization process, DNA quantification was performed. DNA was isolated from 25 mg of dry tissue using the DNeasy® Blood & Tissue Kit (Qiagen, Hilden, Germany), and total DNA content was measured withNanoDrop Lite Plus Spectrophotometer (Thermo Fisher Scientific; Massachusetts, USA)

### 2.6 ECM orthotopic transplantation

Animals in the recipient cohort underwent partial median-lobe hepatectomy (10%). A partial ECM graft was then transplanted into the resected site and sutured with a continuous 6-0 silk suture. Transplanted animals were euthanized 30 days after the procedure.

### 2.7 Recellularization

CTRL ECM, MASH ECM-rat and MASH ECM-human were subjected to prior sterilization with 0.1% (v/v) peracetic acid and 4% ethanol (Proquimios, Rio de Janeiro, Brazil) in ultrapure water. Sterile 1X PBS was used to wash the scaffolds. Sterilization was performed using an orbital shaker at 700 RPM in different cycles, followed by the transfer of the sterilized scaffolds to a non-adherent 48-well-plate previously filled with 1 mL of DMEM-High glucose culture medium supplemented with 10% fetal bovine serum (FBS) and 1% Penicillin-Streptomycin (5,000 Units/mL Penicillin; 5,000 µg/mL Streptomycin). Overnight incubation was performed prior recellularization in order to ensure aseptic handling. For recellularization, HepG2 cells were resuspended in DMEM-High glucose culture medium (433241 Sigma-Aldrich™; Saint Louis, Missouri, USA) supplemented with 10% fetal bovine serum (12657-029 Sigma-Aldrich™) - FBS - and 1% Penicillin-Streptomycin 5,000 Units/mL Penicillin; 5,000 µg/mL Streptomycin (15070063 Sigma-Aldrich™;). Recellularization of each scaffold was performed from cell grafting at a density of 5 × 10⁵ cells in 20 µL - with the scaffold positioned in the center of a cell-repellent surface 96-well-plate (655970 Greiner Bio-One™; Frickenhausen, Germany). Recellularization then consisted of 20 µL of cell suspension grafted onto the top of the scaffold with incubation at 37 °C for 30 min. The step was repeated four times to ensure better cell adhesion to the scaffold. Finally, 140 µL of the complete culture medium were added to each well for overnight incubation at 37 °C. Subsequently, the recellularized scaffolds with HepG2 cells were transferred into a cell-repellent surface 48-well-plate (677970 Greiner Bio-One™) 48-well-plate previously filled with 1 mL of complete medium.

### 2.8 ECM MASH-conditioned medium preparation

Liver specimens derived from MASH rats were obtained during orthotopic liver transplantation performed in recipient animals. The excised tissue was collected during partial hepatectomy and immediately preserved at −80 °C. Following the protocol previously described by Thanapirom et al. (2021) [20], the preserved tissue samples were sectioned into cubic fragments 5mm × 5 mm × 5mm) and individually transferred into microcentrifuge tubes. Cubes (n = 24) were subjected to two sequential thawing steps in a water bath: first at 37 °C for 45 min, followed by immersion in 1× PBS for 15 min at 37 °C. The procedures for decellularization and sterilization were performed in accordance with the protocols previously described for human tissue samples. ECM MASH-rat cubes were then placed in a tissue-treated 6-well-culture plate (Corning 3506) filled with 3 mL complete culture medium/well at 37 °C in an incubator under 5% CO₂. After ten days, the conditioned medium (ECM MASH-rat) was collected. Subsequently, HepG2 cells were seeded onto a 24-well culture-plate (Corning 3524) with sterile coverslips (Figure 6A). Six wells were cultured with ECM MASH-conditioned medium, while the remaining wells received complete growth medium as control group. Media was exchanged every 2-3 days and cells were cultured until 75–80% confluence.

### 2.9 Immunofluorescence staining to trace lipids

BODIPY™ 493/503 (Invitrogen™ D3922; Carlsbad, California) was used as a fluorescent dye for visualization of neutral lipid droplets in HepG2 cell line cultured with ECM MASH-conditioned medium (Qiu B, Simon MC, 2016). The dye was dissolved in dimethyl sulfoxide (DMSO; SigmaAldrich D8418). For staining, the stock solution (10 mM) was diluted in 1X PBS (Nova Biotecnologia 13-30262-05; São Paulo, Brazil) to a final working concentration of 10 µM (Figure 6A). HepG2 cells grown on glass coverslips to 75–80% confluence were incubated with 100 µL of the working solution at room temperature for 30 min, then washed twice with 1X PBS for 5 min each and imaged by fluorescence microscopy using Invitrogen EVOS M5000 (Thermo Fisher Scientific) - maximum excitation/emission wavelength: 493/503 nm. HepG2 cell line cultured with complete growth medium as control were separated into negative - growth medium only - and positive - 30 µM oleic acid - control groups. The positive control group was incubated 30 μM oleic acid (Thermo Scientific AC270290050) for 24 h before staining [21].

### 2.10 Biochemical analysis

Blood samples (500 μL) were collected by cardiac puncture into clot-activator gel microtubes (Vacuplast, São Paulo, Brazil). After collection, samples were centrifuged at 1,400 × g for 10 min, and the resulting serum was stored at −20 °C. Serum levels of albumin (ALB), aspartate aminotransferase (AST), alanine aminotransferase (ALT), high density lipoprotein (HDL), low density lipoprotein (LDL), and triglycerides (TGC) were quantified using a semi-automatic biochemical analyzer (Bio 200, Bioplus, Rio de Janeiro, Brazil). Analyses were performed with the following kits: ALB (Ref. 19), AST (Ref. 109), ALT (Ref. 108) (Labtest, Minas Gerais, Brazil), HDL (Ref. K071-23), LDL (Ref. K088-1), and TGC (Ref. K117-3) (Bioclin, Minas Gerais, Brazil). For blood glucose test, the animals underwent 12 hours fast, then a glucose solution (1g/Kg/ body mass) was administered by gavage, and the blood was collected from the tail 30, 60, 90 and 120 min after gavage

### 2.11 Intracellular Ca²⁺ signalling imaging in recellularized scaffolds

HepG2-recellularized ECM (MASH ECM-rat and human, and CTRL ECM) were maintained in Dulbecco’s Modified Eagle Medium (DMEM) supplemented with 10% fetal bovine serum (FBS) and kept in culture conditions until imaging experiments were performed. Prior to Ca²⁺ imaging, scaffolds were gently rinsed with pre-warmed DMEM, and excess medium was removed. Care was taken to preserve the structural integrity of the recellularized matrices and to avoid detachment of the HepG2 cells. Each scaffold was incubated with 12 µM Fluo-4 AM (Invitrogen, Grand Island, NY, USA) for 30 minutes at 37°C in a humidified 5% CO₂ incubator. After loading, scaffolds were washed with HEPES 1X buffer to remove excess dye and mounted onto glass coverslips for imaging. The recellularized scaffolds were transferred to a custom-built perfusion chamber positioned on the stage of a ZEISS-LSM 880 Airyscan Confocal Microscope. Fluorescence excitation was achieved using a 488-nm laser, and emitted fluorescence was collected using a 515/30-nm bandpass filter. Imaging was performed using a 20× objective lens, and intracellular Ca²⁺ dynamics were monitored in individual HepG2 cells using time-lapse acquisition. Baseline fluorescence was recorded for 30 seconds in HEPES 1X buffer. Scaffolds were then perfused with 50 µM ATP (Sigma Aldrich for 5 minutes to assess Ca²⁺ concentration variations. To data analysis, individual regions of interest corresponding to single HepG2 cells within the scaffold were analyzed with ImageJ, and time-dependent changes in fluorescence were quantified across the entire recording period. Fluorescence intensity values following stimulation (F) were normalized to baseline fluorescence (F₀), and Ca²⁺ responses were expressed as (F/F₀) × 100.

### 2.12 Metabolomics analysis

#### ·2.12.1 Sample Preparation

Liver tissue samples were processed as previously described with minor modifications [22]. Briefly, 200 mg of intact liver tissue were homogenized in phosphate-buffered saline (PBS) at a ratio of 1:3 (w/v). Homogenization was performed using a bead ruptor porcelain bead for three 40 s cycles at 4 m/s, with samples kept on ice between cycles. For metabolite extraction, 100 µL of homogenate were mixed with 1.5 mL of ice-cold isopropanol:methanol (3:1, v/v) containing the internal standards furosemide and *p*-fluoro-DL-phenylalanine (0.2 µg/mL). Samples were vortexed, centrifuged at 15,000 rpm for 15 min at 4 °C, and the supernatant was collected. After a second centrifugation under identical conditions, the supernatant was dried under vacuum and reconstituted in 100 µL of acetonitrile:water (3:1, v/v). To improve metabolite recovery, a second extraction was performed on the remaining protein pellet using methanol:water (1:1, v/v). After vortexing and centrifugation, the supernatant was collected, dried under vacuum, and reconstituted in acetonitrile:water (3:1, v/v). Extracts from both steps were combined, centrifuged to remove particulates, and transferred to LC-MS vials for analysis.

#### ·2.12.2 LC–HRMS Analysis

Untargeted metabolomic analyses were performed using a Vanquish UHPLC system coupled to an Orbitrap Exploris 480 high-resolution mass spectrometer (Thermo Scientific), equipped with a heated electrospray ionization source operating with fast polarity switching between positive and negative modes. Data were acquired over an *m/z* range of 50–1000. Chromatographic separation was achieved on an Acquity HSST3 C18 column (100 × 2.1 mm, 1.8 µm; Waters Corporation). The column temperature was maintained at 35 °C and the autosampler at 4 °C. The flow rate was set to 400 µL/min, and 3 µL of each sample were injected. The mobile phases consisted of ultrapure water (A) and acetonitrile (B), both containing 0.1% formic acid. The gradient elution was performed as follows: 85% A and 15% B at 0 min; the composition was linearly changed to 60% A and 40% B at 3.0 min; then to 5% A and 95% B at 13.0 min, which was maintained until 16.0 min; the system was returned to 85% A and 15% B at 16.5 min and re-equilibrated under these conditions until 20.0 min. The ionization parameters were as follows: spray voltage was set to 3500 V in positive mode and 2500 V in negative mode. The ion transfer tube temperature was set to 325 °C for both modes. Sheath gas flow was set to 50, auxiliary gas to 10, and sweep gas to 3 for both polarities. The maximum spray current was set to 10 in both positive and negative modes. The vaporizer temperature was maintained at 0 °C for both modes. The AGC detection time was set to 20.0 ms in standard mode and 4.0 ms in High Mass (HM) mode. Automatic gain control (AGC) and injection times were optimized to ensure high mass accuracy and sensitivity throughout the analysis.

#### ·2.12.3 LC-MS data processing and statistics

LC-MS raw data files were converted to mzML format using MSConvert. Data Processing, including peak picking, deconvolution, grouping and alignment was performed using MS-DIAL (v4.9.22). A peak list was extracted with a signal-to-noise threshold of 10 and with minimum peak height of 5e5. Missing values were filtered prior to statistical analysis and only features detected in 75% of the samples within at least one experimental group were preserved for downstream analysis. Remaining missing values were assumed to originate from below detection limit and imputed by replacing them with half of the minimum observed value for each feature. Data was normalized using median normalization, scaled using Auto Scaling and log_2_-transformed to reduce systematic variation between samples. Detailed processing parameters are provided in supplementary table 2. Processed data was submitted to MetaboAnalyst (v6.0) for univariate and multivariate statistical analysis and visualization, including fold change, ANOVA, principal component analysis (PCA), partial least squares discriminant analysis (PLS-DA), hierarchical clustering heatmap and violin plots. Metabolites were putatively annotated by MS level 2 by matching exact mass (MS1 tolerance of 0.005 Da; MS2 tolerance of 0.05 Da) and spectra against internal library and NIST database. The identification of shared metabolites and Venn diagram was analyzed using the LC–MS intensity matrix generated in MS-DIAL and analyzed in LibreOffice Calc. Features were considered present when detected (intensity >0) in at least 50% of the samples within each group (≥5/10 for *Tx MASH ECM-CTRL* and ≥3/5 for *Tx MASH ECM-MASH*). Shared metabolites between groups were identified using conditional counting functions. The resulting lists were used to generate a Venn diagram in Intervene to visualize unique and overlapping metabolic features.

### 2.13 Histology analysis

For histological evaluation, biopsies from transplanted livers, decellularized and recellularized ECM were formalin-fixed (24h) and paraffin-embedded. Tissue morphology was assessed using hematoxylin and eosin (H&E) staining (Merck, São Paulo, Brazil), extracellular matrix (ECM) deposition was evaluated using Sirius Red staining (365548; Sigma-Aldrich), and lipid accumulation was assessed using Oil Red O staining (O0625-25G; Sigma-Aldrich). Images were obtained using EVOS microscope (Invitrogen EVOS M5000, Thermo Fisher Scientific).

### 2.14 Immunohistochemistry analysis

For the immunohistochemistry analyses, following deparaffinization and rehydration, the sections were immersed in Tris-buffered saline with 0.1% Tween (TBS-T) for 10 minutes. Then, the slides were submitted to the antigen retrieval step. Antigen retrieval was performed by immersing the slides in pre-heated sodium citrate buffer (pH 6) in a microwave (lower power) at 90-95 °C for 10 min. The slides were then allowed to return to room temperature for approximately 1 hour, followed by a wash in distilled water for 2 min. After that, the slides were exposed to hydrogen peroxide (3%) for 15 min. Excess peroxide was removed with TBS-T for 3 min. Then, the sections were incubated in 2.5 % horse serum albumin for 15 min. Subsequently, the slides were incubated with primary antibodies against desmin (1:50; D1033; Merck) and α-smooth muscle actin (α-SMA) (1:100; Ab7817; Abcam, Cambridge, MA, USA) for 2h at 4 °C. Sections processed ommiting the primary antibodies were used as negative controls. After 2 h, the slides were washed with TBS-T (3 min) and incubated with the Novolink Polymer Detection System (RE7260-CE, Leica Biosystems Newcastle upon Tyne, UK). First, the slides were incubated with post-primary antibody for 30 min (Leica). After that, the excess of post-primary solution was removed with TBS-T for 3 min. Then, the slides were incubated with novocastra polymer solution for 30 min and washed with TBS-T for 3 min. The reaction was developed using 3,3-diaminobenzidine (DAB) (ImmPACT DAB Substrate Kit; REF SK-4105) and subsequently stopped with water. Finally, the sections were counterstained with hematoxylin and mounted with Entellan. Images were acquired using a EVOS microscope (Invitrogen EVOS M5000, Thermo Fisher Scientific).

### 2.15 Scanning electron microscopy

MASH ECM sections were washed three times with 0.1 M sodium cacodylate buffer (pH 7.2) and fixed in 2.5% glutaraldehyde for 24 hours. Subsequently, the samples were washed three times in 0.1 M sodium cacodylate buffer (pH 7.2) and post-fixed for 1 hour in 1% osmium tetroxide (OsO_4_) solution in 0.1 M sodium cacodylate buffer (pH 7.2); then, they were dehydrated in an increasing series of ethanol (30%, 50%, 70%, 90%, and 100%). The fragments were subsequently dried in a critical point dryer (Autosamdri-815; Tousimis, Cambridge, MA, USA), coated with gold in an ion sputtering apparatus (Cressington 108, Watford, UK) and observed under a scanning electron microscope (JEOL-JSM-6390-LV, Akishima, Tokyo, Japan).

### 2.16 Second harmonic generation

Second harmonic generation (SHG) signals recorded at 470 nm were used to capture architectural features of extracellular matrix collagen fibrils through laser excitation at 940 nm, while two-photon excited fluorescence (TPEF) signals recorded were used to visualize tissue structures. Images were acquired at 20× magnification with 512×512 pixel resolution, each image had a dimension of 200 µm × 200 µm.

### 2.17 Quantification of extracellular matrix fiber length and orientation

Quantitative analysis of ECM fibers was performed using ImageJ/Fiji (NIH, USA), following an image-processing workflow adapted from previously validated approaches for automated fiber analysis, including the General Image Fiber Tool (GIFT) methodology adapted fromHuling *et al* [24]. Briefly, fluorescence images were acquired under identical acquisition parameters and analyzed in a blinded manner. Images were converted to grayscale by extracting the red fluorescence channel, corresponding to ECM fiber labeling. Spatial calibration was performed using the embedded scale bar (50 µm), and this calibration was applied uniformly across all images prior to analysis. To reduce background noise while preserving fiber morphology, images were smoothed using a Gaussian blur filter (σ = 1 pixel). Fiber structures were segmented using automatic global thresholding (Otsu method), generating binary images in which ECM fibers were represented as foreground objects. Small, isolated particles and background artifacts were excluded using size-based filtering to ensure that only continuous fibrous structures were retained for quantitative analysis. Binary images were subsequently processed using the Skeletonize function in ImageJ, reducing fibers to one-pixel-wide representations while preserving their topology. This approach is consistent with skeleton-based fiber analysis strategies commonly employed for fibrous networks and is conceptually aligned with the edge-based and morphology-driven framework implemented in the GIFT macro. Quantification of individual fibers was performed using the Analyze Skeleton plugin, which enables the identification of individual fiber segments within the skeletonized image. For each detected fiber, the following parameters were extracted:

a. Fiber length, calculated as the cumulative length of the skeletonized segment and converted from pixels to micrometers using the calibrated image scale;
b. Fiber orientation (angle), determined from the principal axis of each fiber segment and expressed in degrees.

Consistent with the statistical strategy recommended for automated fiber analysis workflows, fiber-level measurements were averaged per image, such that each image represented one independent biological replicate. Group-level comparisons between control and fibrotic (MASH) liver samples were therefore performed using image-level mean values (n = 3 per group), thereby avoiding pseudo-replication arising from the large number of fibers detected within individual images. This adapted ImageJ-based pipeline enables reproducible and unbiased quantification of ECM fiber length and orientation and follows the same core principles of automated, morphology-driven fiber analysis validated by Huling *et al.* for fibrous biomaterials [24].

### 2.18 qRT-PCR

Pro-fibrogenic, tissue remodeling and metabolic targets were assessed by real-time quantitative PCR (RT-qPCR) targeting *Col1a1*, *Timp, Acta2, Acaca, ApoB, PPAR*α*, CPT1*α, *Srebp1c* and *GAPDH*. Total RNA was extracted using the RNeasy Fibrous Tissue Mini Kit (Qiagen, Hilden, Germany). RNA concentration and purity were determined with NanoDrop Lite Plus spectrophotometer (Thermo Fisher Scientific, Wilmington, DE, USA). Complementary DNA (cDNA) was synthesized from 1 µg of total RNA using the High-Capacity Reverse Transcription Kit (Applied Biosystems, Carlsbad, CA, USA) following the manufacturer’s instructions. Quantitative PCR reactions were performed on a QuantStudio™ 3 Real-Time PCR System (Applied Biosystems). Each reaction was conducted in duplicate with a final volume of 15 µL, containing 10 ng of cDNA, 7.5 µL of 2×Power SYBR Green Master Mix (Applied Biosystems), RNase-free water and 0.5 µL each of forward and reverse primers (10 µM). Thermal cycling conditions consisted of an initial incubation at 50 °C for 2 min, followed by enzyme activation at 95 °C for 10 min, and 40 cycles of denaturation at 95 °C for 15 s, primer annealing at gene-specific temperatures (listed in Supplementary Table S1) for 30 s, and extension at 72 °C for 30s. Gene expression levels were normalized to GAPDH as the endogenous control. Relative transcript abundance was calculated using the 2 -(ΔΔCt)) method, where ΔΔCt was determined by comparing experimental samples with control livers. Data are presented on a logarithmic scale (base 10). Primer sequences for the analyzed genes are listed in Supplementary Table S1.

### 2.19 -Statistical analysis

Statistical analyses were performed through GraphPad Prism 10 (GraphPad Software, La Jolla, CA, USA). Descriptive data are expressed as means ± standard deviation (SD). Group comparisons were assessed using paired or unpaired student’s t-tests with Welch’s correction. One-way ANOVA followed by Tukey’s post hoc test, or two-way ANOVA followed by Šídák’s multiple comparisons test were applied. Statistical significance was set at p < 0.05.

## 3. Results

### High fat diet + CCl_4_ induces steatohepatitis in rats

The first set of experiments was performed to obtain rats with steatohepatitis. Wistar rats subjected to a multiple hit MASH induction (Figure 2A) showed differences in body weight when compared to the control group (Figure 2B). Biochemical analysis showed that rats submitted to steatohepatitis protocol significantly elevated serum ALB, ALT, and AST (Figure 2C). In addition, a significant increase in TGC, HDL, LDL, were observed similar to the biochemical alterations described in human liver steatohepatitis elsewhere (Figure 2C). No differences were observed in serum glucose levels between groups (Figure 2D).

**Figure 2.**
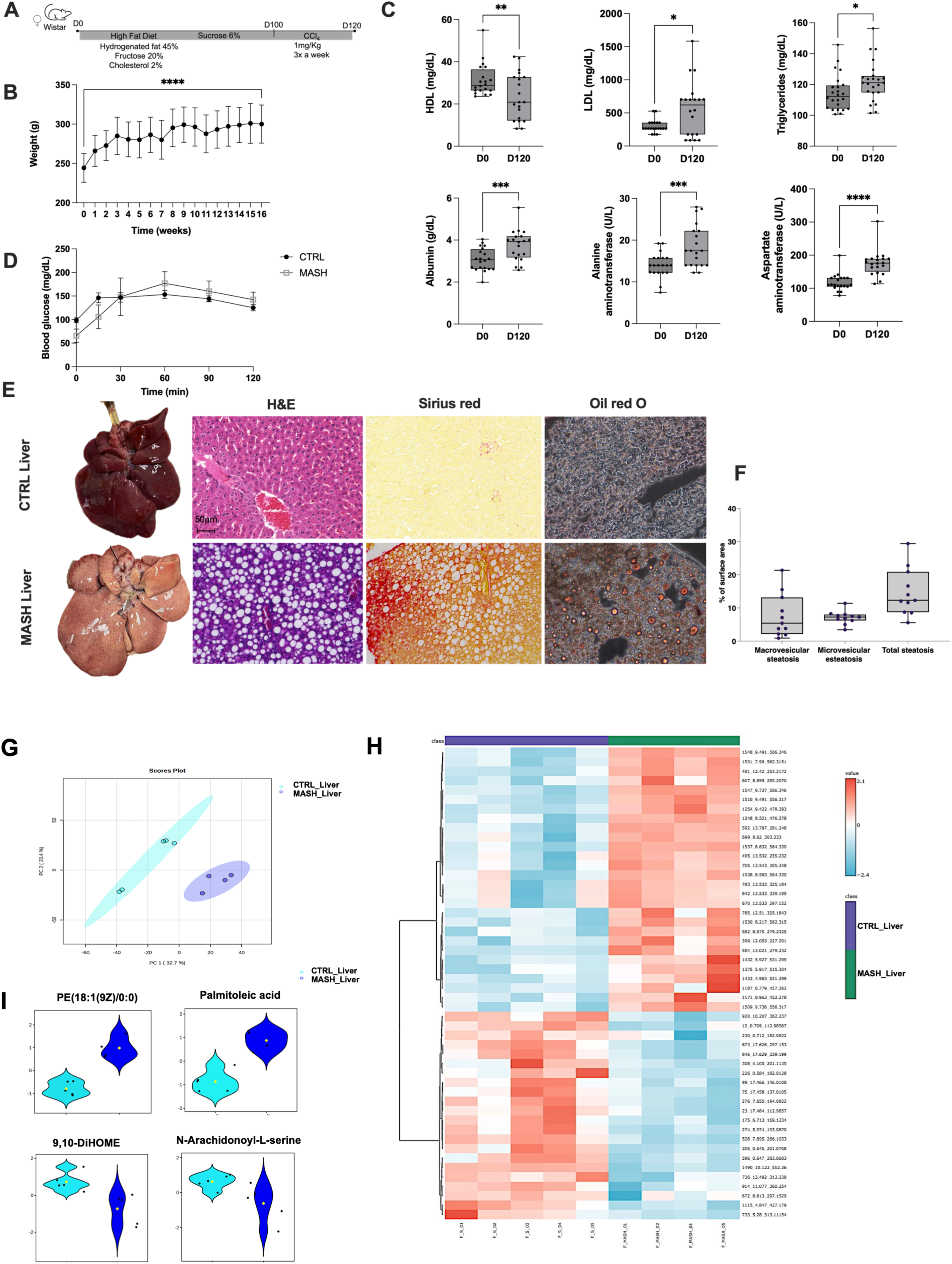
Characterization of MASH rat livers. (A) MASH livers were obtained after high-fat diet administration in Wistar rats for 120 days. (B) Body weights of Wistar rats during the MASH induction protocol. (C) HDL, LDL, TGC, Albumin, ALT and AST levels in the serum of Wistar rats before and after MASH induction. (D) Blood glucose levels in healthy and MASH rats. (E) Macroscopic and microscopic views of healthy (control) and MASH livers. Hematoxylin and Eosin, Sirius red, and Oil red O-stained sections from control and MASH livers. Scale bars: 50 μm. (F) Percentage of microvesicular, macrovesicular, and total steatosis in MASH livers. (G) PCA score plot showing the separation between CTRL_Liver and MASH_Liver samples based on metabolomic profiles. (H) Heatmap of discriminant metabolites illustrating the hierarchical clustering of samples and metabolites (red: higher abundance; blue: lower abundance). (I) Bean plots of representative metabolites (PE(18:1(9Z)/0:0), palmitoleic acid, 9,10-DiHOME, and N-arachidonoyl-L-serine) highlighting differences in relative abundance between the CTRL_Liver and MASH_Liver groups; dots represent individual samples, and diamonds indicate group means. The results are presented as mean ± SD. Statistical significance between the control and MASH groups was determined using the Student’s t-te,st where *p <0.05, **p <0.01, ***p <0.001, and ****p <0.0001 (n= 20). CTRL, Control group; MASH, metabolic dysfunction-associated steatohepatitis.

In contrast with control liver, macroscopic analysis showed that our protocol generates yellowish livers that were lighter in color with white regions suggestive of lipid deposition (Figure 2E). While hepatocytes aligned to sinusoids and absence of steatosis was observed in the control liver tissue, histological analysis of MASH livers stained with H&E confirmed inflammation, hepatocyte ballooning, macrovesicular and microvesicular steatosis (Figure 2E). After quantitative analysis we confirmed that most of the liver surface was occupied by macrovesicular steatosis (Figure 2F). This intense lipid deposition was confirmed by Oil red O staining (Figure 2E). In addition to that, Sirius red staining confirmed the presence of fibrosis distributed throughout the liver parenchyma (Figure 2E).

A comprehensive untargeted metabolomic analysis was performed using liquid chromatography coupled to high-resolution mass spectrometry (LC-HRMS) in both positive and negative electrospray ionization modes (ESI⁺/ESI⁻) to characterize metabolic alterations associated with MASH. Multivariate statistical analyses were applied to evaluate global metabolic differences between CTRL and MASH liver samples. Partial Component Analysis (PCA) (Figure 2G) revealed a clear separation between CTRL Liver and MASH Liver groups, indicating a robust disease-associated metabolic signature. Component 1 (32.7%) and Component 2 (23.4%) jointly explained a substantial proportion of the variance related to disease status. Model performance metrics demonstrated high classification accuracy (1.0), strong goodness-of-fit (R² up to 0.97), and satisfactory predictive ability (Q² ≈ 0.78–0.83). Permutation testing (1,000 permutations) confirmed the statistical robustness of the model, with the observed class separation significantly exceeding that obtained from randomized labels, minimizing the risk of overfitting. Hierarchical clustering analysis of discriminant *m/z* features further supported these findings, revealing distinct metabolomic profiles between CTRL Liver and MASH Liver groups (Figure 2H) based on combined criteria of statistical significance (p < 0.05) and magnitude of change (|log₂FC| ≥ 1), this approach allowed prioritizing biologically relevant metabolites, reducing the impact of variations of low pathophysiological relevance. The heatmap analysis (Figure 2H) highlights a predominant cluster of metabolites increased in MASH Liver samples, largely composed of lipid-related species such as phospholipids, lysophospholipids, fatty acids, and bioactive lipid mediators. The coordinated accumulation of these species is consistent with dysregulation of hepatic lipid metabolism, remodeling of cell membranes, lipotoxicity, and activation of inflammatory pathways—processes central to the pathogenesis of MASH. Consistent with the heatmap analysis, the beanplot analysis (Figure 2I) highlights some specific lipids classes, as PE (18:1(9Z)/0:0) and palmitoleic acid increased in MASH samples, supporting enhanced phospholipid turnover and activation of de novo lipogenesis. In contrast, other lipid-derived mediators, including 9,10-DiHOME and *N*-arachidonoyl-*L*-serine, are significantly reduced in MASH samples. 9,10-DiHOME is an oxylipin derived from linoleic acid metabolism and participates in oxidative lipid signalling, whereas N-arachidonoyl-L-serine belongs to the family of endocannabinoid-related lipids involved in inflammatory and metabolic regulation. Their depletion suggests a disruption of lipid mediator signalling pathways that normally contribute to metabolic homeostasis and resolution of inflammation. These metabolites represent distinct lipid classes, indicating broad alterations in hepatic lipid metabolism associated with disease status.

### MASH promotes biochemical and ultrastructural alterations in ECM

MASH livers were submitted to decellularization using a detergent-based perfusion protocol adapted from our previous report (Dias et al., 2023) [7]. While decellularization of healthy livers generated a translucent ALS, the decellularization of MASH livers originated a dense, opaque and white-colored MASH ECM (Figure 3A). Weight quantification analysis showed that MASH ECM is significantly heavier than control ALS (Figure 3B). In addition to macroscopic and weight differences, histological analysis also showed important differences between control and MASH ECM (Figure 3A). Histological sections stained with H&E showed that MASH ECM is composed of a dense ECM, while a less abundant ECM is present in control ECM (Figure 3A). Sirius red staining showed that MASH ECM is composed of an abundant collagen content. In contrast, control ECM showed less intense Sirius red staining (Figure 3A). SEM analysis also revealed important differences between control and MASH ECM (Figure 3C). While an ECM with preserved collagen fibers, following general micro and ultra-structure tissue preservation was observed in the control ECM, MASH ECM showed a dense ECM structure, with a different arrangement and conservation of the three-dimensional liver ECM (Figure 3C). These differences were also explored by SHG analysis (Figure 3C). A distinct pattern of organization and disposition of collagen fibers were observed in MASH ECM when compared to the control ECM (Figure 3C). The collagen network exhibited bundling in MASH ECM whereas the network appeared to be more organized with thinner fibres in control ECM. Significant differences were observed when the fiber length was compared between the groups (Figure 3D). No differences were observed regarding fiber angle when MASH ECM and CTRL ECM were compared (Figure 3E).

**Figure 3.**
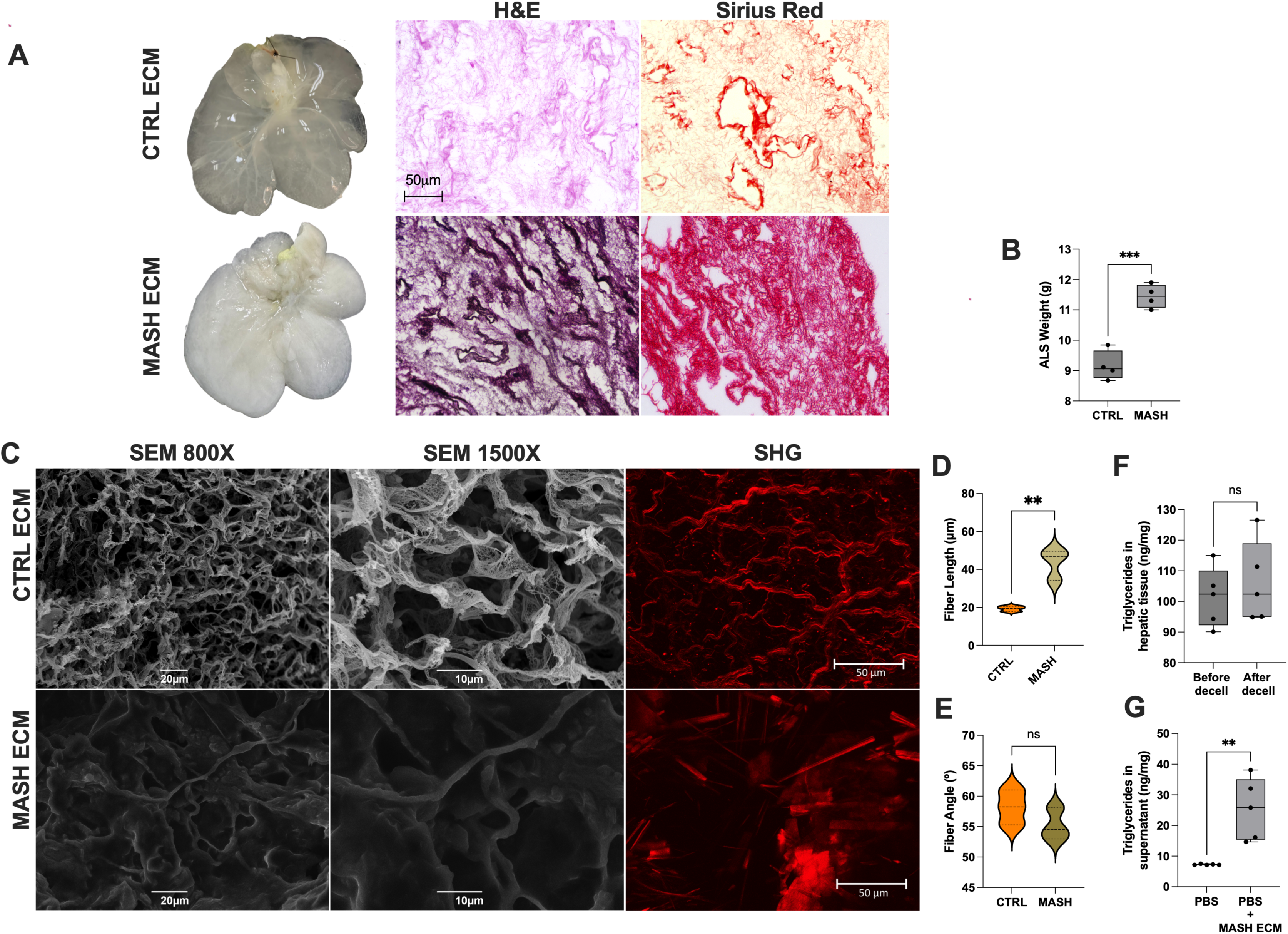
Characterization of decellularized MASH ECM. (A) Macroscopic and microscopic views of decellularized control ECM and MASH ECM obtained after decellularization. Hematoxylin and Eosin and Sirius red-stained sections from control and MASH decellularized ECM. Scale bars: 50 μm. (B) Weights of decellularized control and MASH ECM (n= 4). (C) Ultrastructural characterization of control and MASH ECM. Scanning electron microscopy images at 800x and 1500x magnification obtained from control and MASH ECM. Scale bars: 20 and 10 μm, respectively. Second Harmonic Generation (SHG) images of control and MASH ECM. Scale bars: 50 μm. (D) Quantification of fiber length, expressed as the mean fiber length per image (n = three images per group). (E) Quantification of fiber orientation (angle), calculated as the mean fiber angle per image (n = 3 images per group). Statistical comparisons were performed using the image-level averages to avoid pseudo-replication. (F) Triglyceride content in the hepatic tissue before and after decellularization (n= 5). (G) Triglyceride content in the supernatant of MASH ECM, where PBS was used as a control (n= 5). The results are represented as mean ± SD. Statistical significance between the groups was determined using the Student’s t-test where *p <0.05, **p <0.01, ***p <0.001, and ****p <0.0001.

### Lipids are present in the MASH ECM after decellularization

Given the nature of MASH disease, highlighted by lipid accumulation in the liver tissue, our next question focused on evaluating the presence of triglycerides in the MASH ECM. To directly test the presence of triglycerides in the ECM, MASH ECM was subjected to triglyceride quantification after decellularization. The TGC assay showed that the MASH ECM contained triglyceride content at similar levels found in the MASH liver before decellularization (106.0±13.31 ng/mg; 101.4±9.72 ng/mg, respectively) (Figure 3F). Considering this result, we also evaluated the presence of triglyceride content in the supernatant (PBS) where MASH ECM was stored previously to the transplantation procedure to further explore the possibility that the triglyceride embedded in the MASH ECM could be released in the storage media. TGC assay analysis confirmed the presence of triglyceride content in the supernatant where MASH ECM was stored for 24h and the data demonstrated a significant release of triglyceride content in the media of MASH ECM when compared to control media (7.25±0.16 ng/mg; 25.35±10.09 ng/mg, respectively) (Figure 3G).

### MASH ECM maintain steatohepatitis after transplantation

From a translational perspective and to evaluate the use of an ECM derived from MASH liver in transplantation we promoted a MASH-ECM partial orthotopic transplantation in recipient MASH rats. Exploratory laparotomy after euthanasia 30 days after transplantation revealed the assembly of a soft textured, yellowish colored tissue connected to the recipient rat median lobe (Figure 4A). Histological analyses were performed to investigate whether MASH-ECM were able to be recellularized after transplantation in MASH recipients. Histological sections stained with H&E revealed that MASH-ECM was repopulated by a large number of cells, including mononuclear cells (Figure 4A). Notably, we observed that the recellularization was followed by the detection of vesicular structures similar to those found in the MASH liver before transplantation suggesting the presence of macrovesicular steatosis into MASH-ECM 30 days after transplantation (Figure 4A). Sirius red staining revealed a colocalization between steatosis and collagen fibers (Figure 4A).

**Figure 4.**
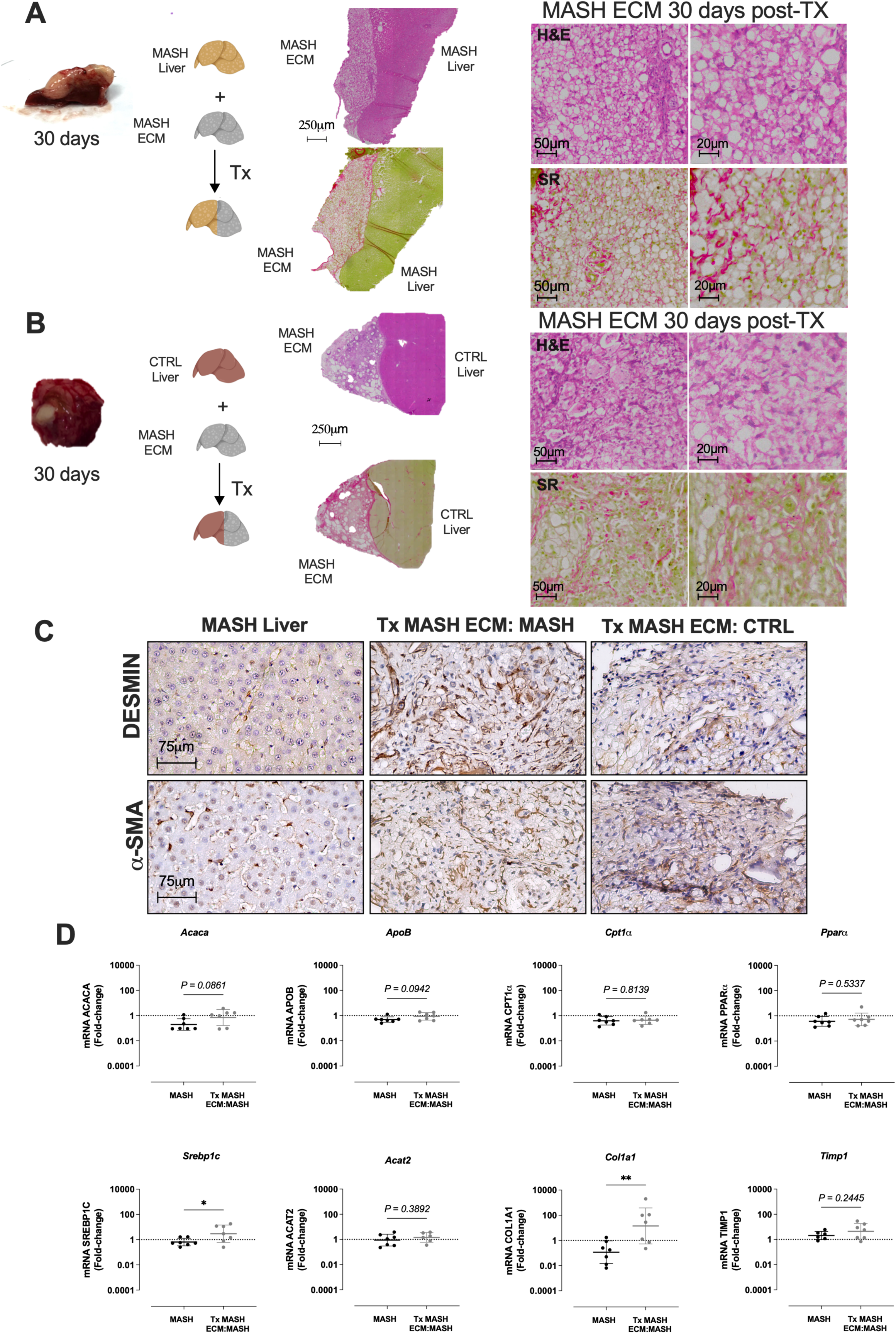
Microscopic, macroscopic, and molecular analyses of MASH ECM after transplantation in healthy and MASH recipients. (A) Macroscopic and microscopic views of biopsies from the MASH ECM 30 days after transplantation in MASH recipients. Histological sections (20x, 40x magnification) were stained with hematoxylin and eosin and Sirius red. Scale bars: 50 µm and 20 ∝μ, respectively. (B) Macroscopic and microscopic views of biopsies from MASH ECM 30 days after transplantation in control recipients. Histological sections (20x, 40x magnification) were stained with hematoxylin and eosin and Sirius red. Scale bars: 50 µm and 20 ∝μ, respectively. (C) Immunohistochemical staining of desmin and α-SMA – positive cells in MASH liver (positive control), MASH ECM transplanted in MASH recipients (Tx MASH ECM: MASH), and MASH ECM transplanted in control recipients (Tx MASH ECM: CTRL) 30 days after transplantation. Scale bars: 75 μm. (D) Expression of genes linked to lipid metabolism and tissue fibrosis, *Acaca, ApoB, CPT1α, PPARα, Srebp1c, Acat2, Col1a1, and Timp1,* in the MASH liver and MASH ECM transplanted in MASH recipients. Gene expression levels were normalized to GAPDH as the endogenous control, and the data are presented on a logarithmic scale (base 10). Statistical significance between groups was determined using an unpaired Student’s t-test with Welch’s correction, where *p <0.05 and **p <0.01 (n= 7).

Given these results, our next question was to determine whether the steatosis maintenance could be promoted by the MASH ECM. To directly test the effect of a MASH ECM, we performed MASH ECM transplantation in recipient healthy rats. Histological analysis performed 30 days after transplantation revealed similar findings even when MASH ECM is transplanted in recipient healthy rats (Figure 4B). H&E-stained sections confirmed the presence of vesicular structures suggesting the presence of macrovesicular steatosis into MASH ECM 30 days after transplantation in healthy recipient rats (Figure 4B). Colocalization between steatosis and collagen fibers was also observed (Figure 4B). As we observed ECM deposition in both transplanted MASH ECM, we performed immunohistochemistry analysis to confirm the presence of hepatic stellate cells and myofibroblasts into the MASH ECM after transplantation. We observed cells expressing both desmin and alpha smooth muscle actin (α-SMA) in MASH ECM after transplantation in MASH and CTRL recipients. MASH recipient liver was used as a positive control (Figure 4C).

In addition to histological evaluation, biochemical analysis was performed to investigate whether the MASH ECM could impact liver metabolism, function and injury parameters (Supplementary material). HDL levels significantly increased, while LDL levels significantly decreased 30 days after transplantation. No significant differences were observed in triglycerides content before and after transplantation (Supplementary material). In addition, MASH ECM transplantation significantly decreased albumin levels and did not promote changes in ALT and AST levels (Supplementary material).

### MASH ECM maintains the expression of genes involved in lipid metabolism and induces fibrosis

In light of findings suggesting that MASH ECM promotes steatohepatitis maintenance after transplantation, we investigated this aspect at the molecular level. To address this issue, the mRNA expression levels for lipid metabolism and pro-fibrogenic and tissue remodelling genes were also investigated. Our data showed no significant differences in *Acaca, Apob, Cpt1a, Ppara, Acat2 and Timp1* gene expression when MASH livers and MASH ECM transplanted in MASH recipients were compared. However, a significant increase in *Screbp1c,* which plays a central role in controlling the expression of genes involved in de novo lipogenesis, and *Col1a1,* which is involved in ECM production and fibrosis, was observed (Figure 3D), confirming our previously obtained data from histological analysis.

### MASH ECM acquires a metabolic profile similar to that of the MASH liver after transplantation

To explore whether de MASH ECM can impact the metabolic profile of resident cells and promote steatohepatitis, we collected ECM MASH transplanted into MASH recipient rats (Tx MASH ECM-MASH) for metabolomic analysis (Figure 5). The metabolomic data were compared with MASH ECM transplanted into healthy recipient rats (Tx MASH ECM-CTRL).

**Figure 5.**
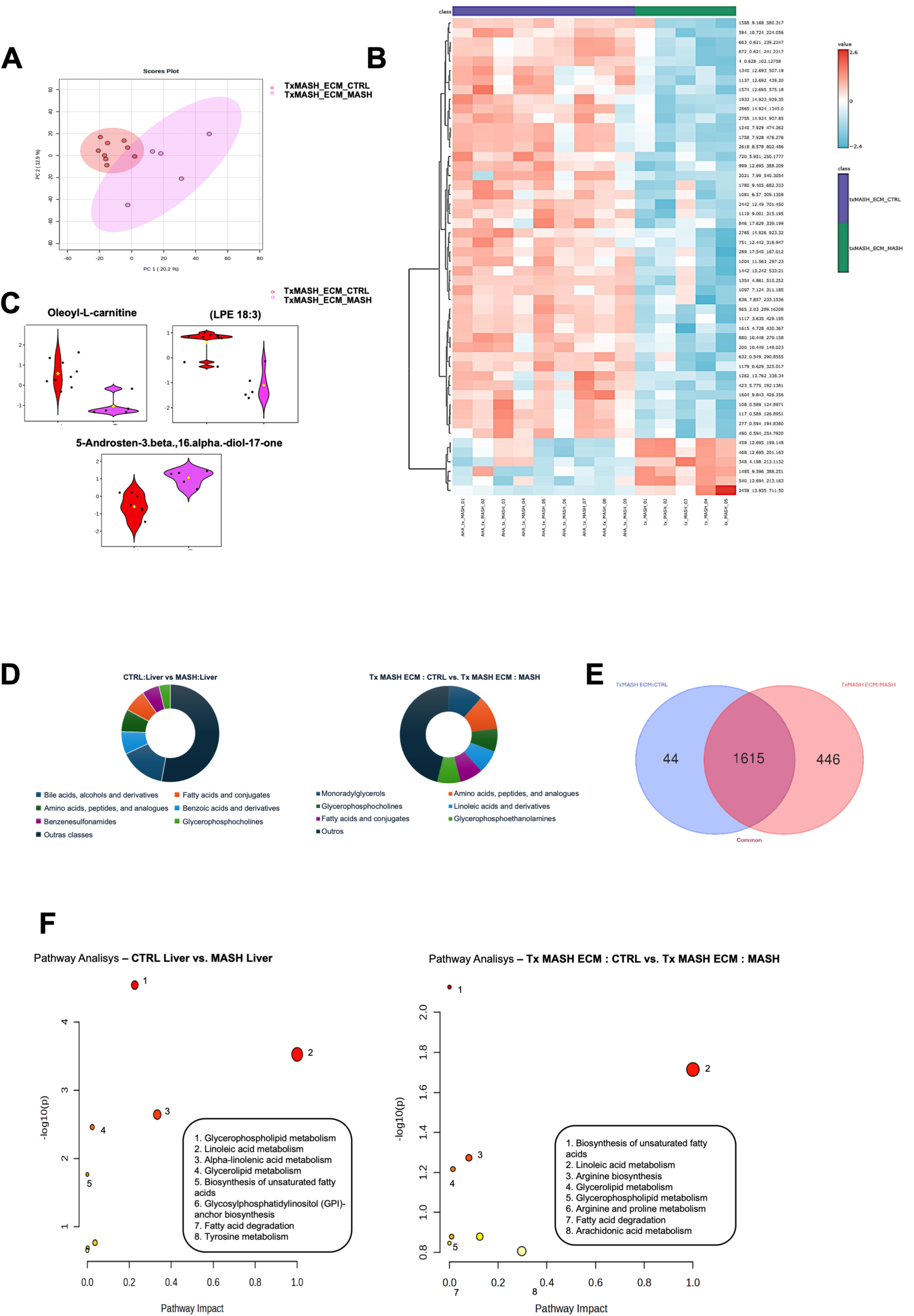
Metabolomic similarities and differences between MASH ECM transplanted in MASH and Control recipients. (A) PCA score plot showing the separation between Tx MASH ECM-CTRL and Tx MASH ECM-MASH Liver samples based on metabolomic profiles. (B) Heatmap of discriminant metabolites illustrating the hierarchical clustering of samples and metabolites (red: higher abundance; blue: lower abundance). (C) Bean plots of representative metabolites (oleoyl-L-carnitine, LPE 18:3 and 5-Androsten-3. beta.,16.alpha.-diol-17-one) highlighting differences in relative abundance between Tx MASH ECM-CTRL and Tx MASH ECM-MASH; dots represent individual samples, and diamonds indicate group means. (D) Donut charts showing the chemical subclass distribution of VIP metabolites identified in the comparisons between CTRL Liver and MASH Liver and Tx MASH ECM-CTRL and Tx MASH ECM-MASH. Each segment represents the relative proportion of metabolites within each chemical subclass among the selected VIP features. (E) Venn diagram showing the overlap of metabolic features between Tx MASH ECM-CTRL and Tx MASH ECM-MASH samples. A substantial number of shared metabolites (1615) indicates a largely conserved metabolic composition of the transplanted MASH ECM, while a smaller subset of unique metabolites suggests selective metabolic modulation by the hepatic microenvironment. (F) Comparative pathway enrichment analysis between control and MASH liver tissue and transplanted ECM conditions.

First, we attempted to investigate the metabolomic profile of MASH ECM to understand whether it can be different or similar to that of the recipient liver. We observed that MASH ECM acquired a similar metabolic profile even when transplanted in healthy or MASH recipient rats (Figure 5). PCA of transplanted MASH ECM samples showed a less pronounced separation (Figure 5A) when compared to PCA of CTRL livers x MASH livers (see Figure 2G). Tx MASH ECM-CTRL and Tx MASH ECM-MASH samples clustered separately along PC1 (20.2%) and PC2 (12.9%), although partial overlap was observed. The MASH ECM transplantation analysis revealed that the host microenvironment partially influences the molecular profile of the transplanted scaffold; however, some metabolites are common, independent of whether MASH ECM is transplanted in healthy or MASH recipients. The greater dispersion observed in the Tx MASH ECM-MASH group suggests that the diseased hepatic microenvironment promotes heterogeneous ECM remodelling, potentially reflecting differences in fibrosis progression, inflammatory activity, or stellate cell activation.

Hierarchical clustering analysis of the transplanted MASH ECM samples (Figure 5B) revealed distinct molecular patterns between Tx MASH ECM-CTRL and Tx MASH ECM-MASH groups. Most features displayed higher relative abundance in Tx MASH ECM-CTRL samples and reduced levels in Tx MASH ECM-MASH samples, suggesting a global reduction of several ECM-associated molecules in the MASH hepatic microenvironment. Conversely, a smaller cluster of features exhibited the opposite trend, with increased abundance in Tx MASH ECM-MASH samples, potentially reflecting ECM remodeling processes induced by the diseased liver environment. The beanplot analysis (Figure 5C) of selected discriminant metabolites between Tx MASH ECM-MASH and Tx MASH ECM-CTRL revealed distinct metabolic signatures associated with the hepatic microenvironment in which the MASH ECM scaffold was transplanted. Notably, Oleoyl-*L*-carnitine and lysophosphatidylethanolamine (LPE 18:3) were more abundant in MASH ECM samples recovered from control livers, whereas 5-Androsten-3β,16α-diol-17-one was elevated in MASH ECM transplanted into MASH livers. Oleoyl-*L*-carnitine is an acylcarnitine involved in mitochondrial fatty acid transport and β-oxidation, and its reduced levels in the Tx MASH ECM-MASH group may reflect impaired lipid oxidation and mitochondrial dysfunction, metabolic features commonly associated with MASH. Similarly, the decreased abundance of LPE (18:3), a lysophospholipid generated during membrane phospholipid remodeling, may indicate altered lipid turnover and inflammatory signaling within the diseased hepatic microenvironment. In contrast, the increase in the steroid-derived metabolite 5-Androsten-3β,16α-diol-17-one in ECM exposed to MASH livers suggests alterations in steroid metabolism, consistent with the known role of the liver in steroid biotransformation and the endocrine disturbances reported in metabolic liver disease. Together, these findings support the concept that the MASH hepatic microenvironment actively reprograms the metabolic composition of transplanted ECM scaffolds, reflecting the systemic metabolic disturbances characteristic of steatohepatitis.

After these observations, we explored the similarities and differences between CTRL Liver, MASH Liver, Tx MASH ECM-CTRL and Tx MASH ECM-MASH by analysing the chemical nature of metabolites found in metabolomic analysis. The donut charts (Figure 5D) illustrate the chemical subclass distribution of VIP metabolites identified in the comparisons CTRL Liver vs MASH Liver and Tx MASH ECM- CTRL vs Tx MASH ECM-MASH. In the CTRL Liver vs MASH Liver comparison, the metabolite profile was strongly dominated by bile acids, alcohols and derivatives, indicating that metabolites associated with bile acid metabolism represent the main discriminant class between healthy and MASH liver samples. Other subclasses were present at lower proportions, including fatty acids and conjugates, amino acids, benzoic acid derivatives, benzenesulfonamides, and glycerophosphocholines. In contrast, the Tx MASH ECM-CTRL vs Tx MASH ECM-MASH comparison exhibited a more heterogeneous distribution of metabolite classes, with notable contributions from monoradylglycerols, amino acids, glycerophosphocholines, fatty acids and conjugates, linoleic acid derivatives, and glycerophosphoethanolamines. This pattern suggests that lipid-related metabolites and phospholipid subclasses play an important role in the metabolic profile observed in the transplanted MASH ECM, potentially reflecting lipid remodeling and metabolic signaling processes within the MASH hepatic microenvironment. In addition to the differences, we also observed similarities between Tx MASH ECM-CTRL and Tx MASH ECM-MASH groups. The donut charts showed that the chemical class distribution revealed a highly similar metabolic composition between Tx MASH ECM-CTRL and Tx MASH ECM-MASH, with both groups predominantly enriched in lipid-related metabolites such as monoradylglycerols, glycerophosphocholines and fatty acid derivatives. This profile becomes more evident when we analyze the Venn diagram of the comparison between Tx MASH ECM-CTRL and Tx MASH ECM-MASH. A total of 1,615 metabolites were shared between the two conditions, whereas only 44 and 446 metabolites were uniquely detected by the groups (Figure 5E). This large intersection highlights a strong similarity in the metabolic composition of the transplanted ECM regardless of the hepatic microenvironment. Such overlap suggests that the MASH ECM retains a conserved biochemical signature, likely reflecting its structural components and lipid-associated metabolites.

Then, we investigated the lipid-related metabolic pathways in MASH ECM transplanted in healthy and MASH recipients. Pathway enrichment analysis revealed that lipid-related metabolic pathways were prominently altered in both comparisons (Figure 5F). In CTRL Liver vs MASH Liver samples, significant pathways included glycerophospholipid metabolism, linoleic acid metabolism, glycerolipid metabolism, and biosynthesis of unsaturated fatty acids, highlighting profound alterations in hepatic lipid metabolism associated with MASH. Interestingly, several of these pathways were also enriched in the ECM transplantation analysis (Tx MASH ECM-CTRL vs Tx MASH ECM-MASH), suggesting that the metabolic environment of the MASH liver influences the biochemical composition of the transplanted ECM. In addition to lipid metabolism, ECM-specific enrichment of arginine and proline metabolism and arachidonic acid metabolism was observed, pathways closely related to extracellular matrix remodeling, inflammatory signaling, and fibrogenesis.

### MASH ECM promotes lipid accumulation in vitro

We next evaluated the impact of MASH ECM in vitro. We first submitted MASH ECM to culture with a basal medium for 10 days (Figure 6A). Lipids droplets were observed in the medium during all the culture time (Figure 6B). The medium (conditioned medium) was collected every other day. Then, we cultured HepG2 cells with the medium collected from MASH ECM culture to investigate whether ECM-associated factors and lipids associated with MASH ECM could solely induce lipid accumulation in HepG2 (Figure 6C). Next, we used both phase-contrast imaging and fluorescence imaging where boron-dipyrromethene (Bodipy) was used to stain lipid droplets. Stained HepG2 cells revealed lipid accumulation after MASH ECM conditioned medium culturing similarly to the positive control where HepG2 cells were cultured with oleic acid (positive control). In contrast, less intense staining was observed in HepG2 cells cultured with control medium (Figure 6C). These qualitative observations were confirmed by a quantitative analysis of fluorescence intensity (Figure 6D). Then MASH ECM was recellularized with HepG2 cells to investigate whether the MASH ECM could promote lipid accumulation in vitro when cells directly interact with the ECM (Figure 6E). H&E-stained sections showed that HepG2 cells also accumulated lipids after culture in MASH ECM for 10 days (Figure 6F).

**Figure 6.**
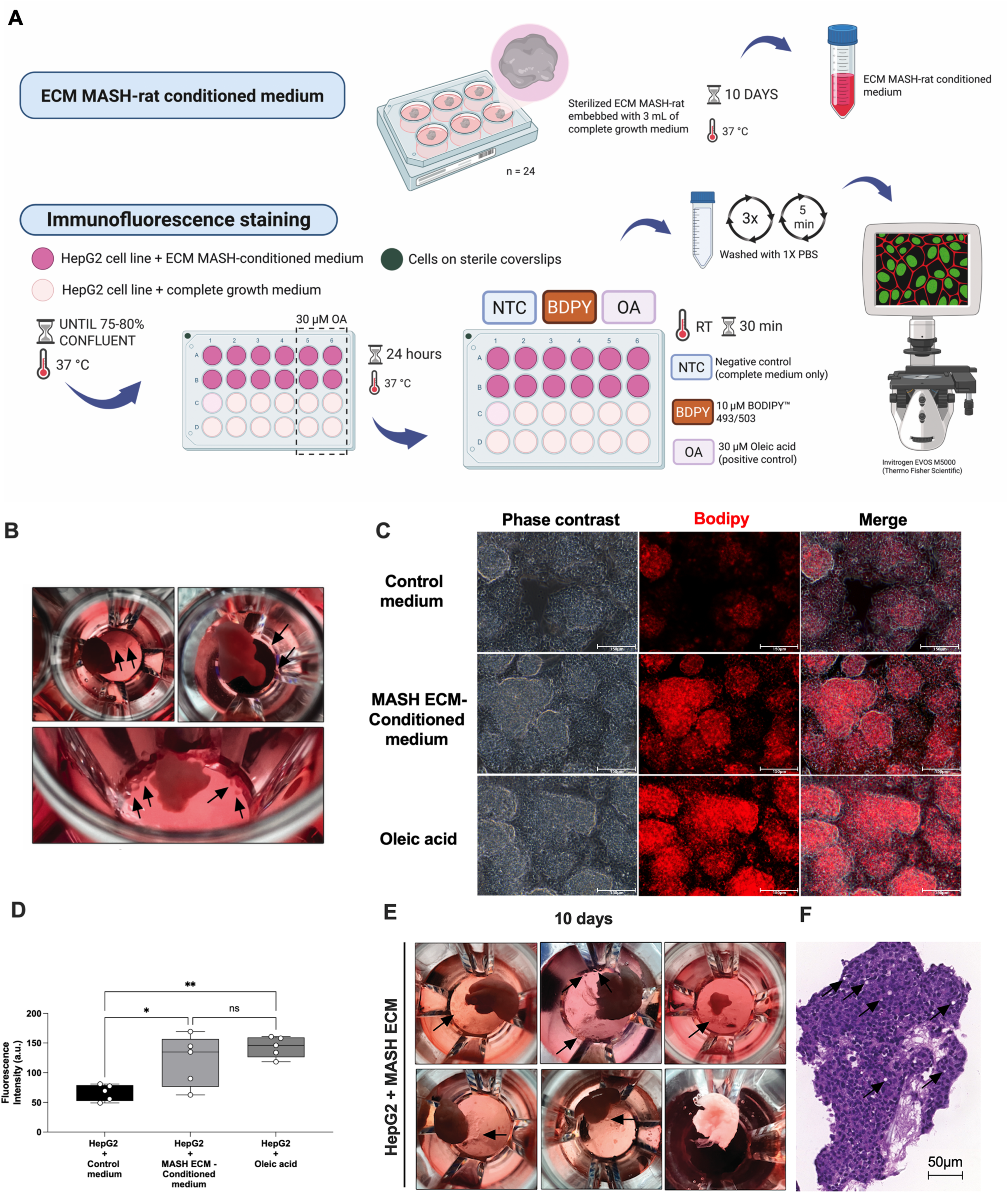
In vitro analysis of MASH ECM and MASH ECM-associated factors on lipid accumulation by HepG2 cells. (A) Experimental design used to investigate the impact of MASH ECM-associated factors on HepG2 cells in vitro. (B) Macroscopic view of MASH ECM in culture without cells. Black arrows indicate lipid droplets in the wells in which MASH ECM was cultured for 10 days. (C) Phase-contrast and Bodipy-stained HepG2 cells after culture with control medium, conditioned medium derived from MASH ECM, and oleic acid (positive control). Scale bars: 150 μm. (D) Fluorescence intensity of Bodipy-stained HepG2 cells after culture with control medium, conditioned medium derived from MASH ECM, and oleic acid. (E) MASH ECM after recellularization with HepG2 cells for 10 days in culture. Black arrows indicate lipid droplets. (F) Hematoxylin and eosin-stained section from MASH ECM recellularized with HepG2 cells. Black arrows indicate cells with lipid accumulation. Scale bars: 50 μm.

### MASH ECM reduces calcium signalling amplitude after recellularization

After in vivo and in vitro exploration, MASH ECM were also explored in ex vivo experiments. MASH ECM cubes were recellularized with HepG2 cells and then submitted to intracellular Ca²⁺ signaling analysis to investigate whether MASH ECM environment could impact on cell physiology properties (Figure 7). Time-lapse confocal images of Fluo-4–loaded HepG2 cells within MASH ECM or CTRL ECM showed that in both conditions cells were able to be responsive to ATP stimulation (Figure 7A). However, the quantification of ATP-evoked Ca²⁺ signals revealed that HepG2 cells cultured in MASH ECM were less responsive to ATP than observed in CTRL ECM (Figure 7B). In addition to that, we also observed that the calcium signalling amplitude was significantly reduced in MASH ECM recellularized with HepG2 when compared to recellularized CTRL ECM (Figure 7C). Moreover, in MASH ECM a change in the pattern of the Ca²⁺ transient was also observed, with a high number of cells responding in a wave manner (Figure 7D). Immunohistochemistry analysis for intracellular calcium channel isoforms 1, 2 e 3 (ITPR1, 2 e 3) were also evaluated (Figure 7E). All isoforms were identified in HepG2 cultured in both CTRL and MASH ECM. However, a more intense staining was observed for ITPR3 isoform in HepG2 cultured in MASH ECM (Figure 7E).

**Figure 7.**
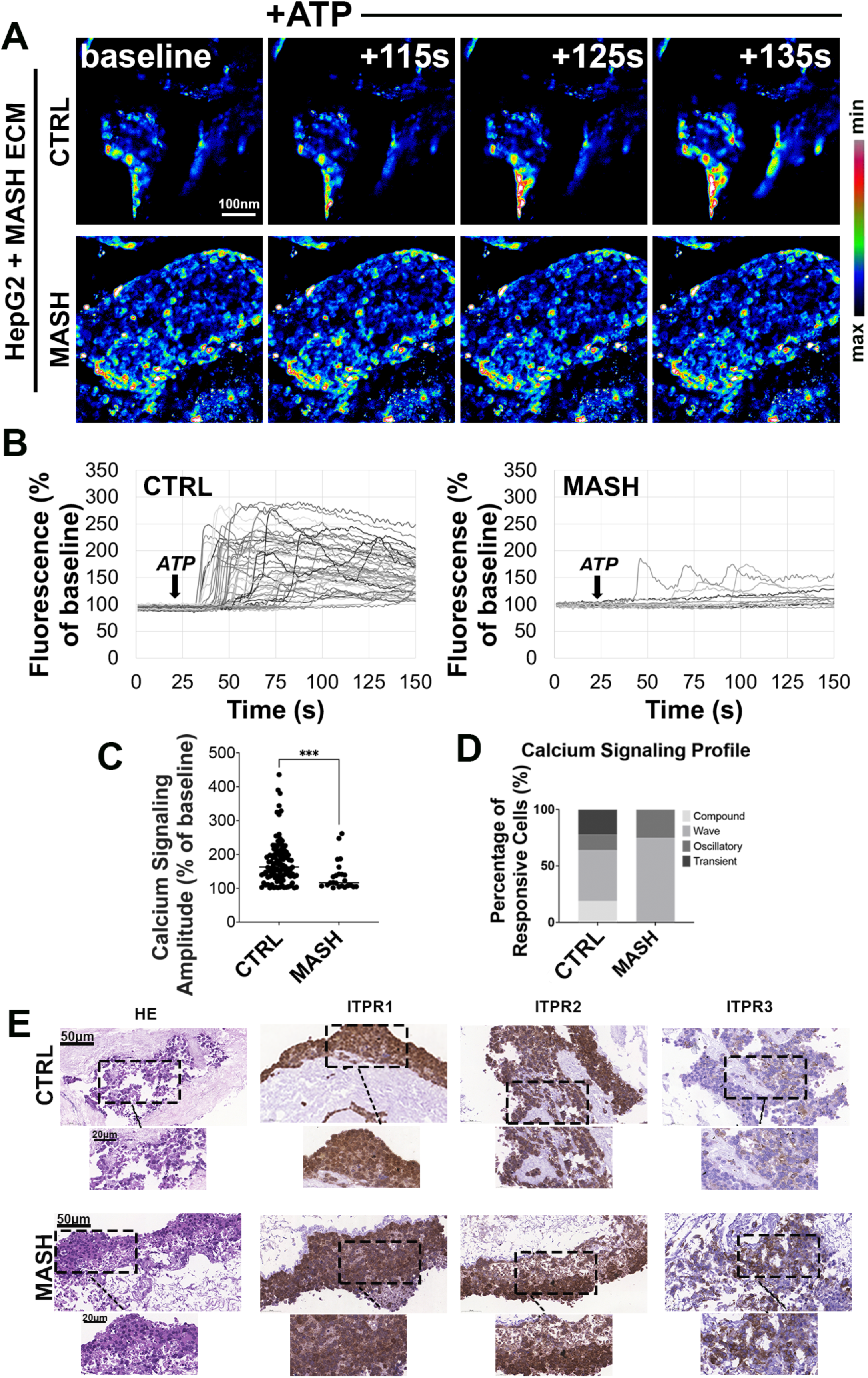
Ca²⁺ dynamics in HepG2-recellularized MASH ECM derived from rats. (A) Representative time-lapse confocal images of Fluo-4–loaded HepG2 cells within rat decellularized scaffolds at baseline and following ATP stimulation. The images show fluorescence frames at +115, +125, and +135 s after the onset of stimulation. The pseudocolor scale (right) indicates the minimum to maximum intracellular Ca²⁺ fluorescence intensity. Scale bar: 100 nm. (B) Quantification of ATP-evoked Ca²⁺ signals in HepG2 cells, with individual traces representing single-cell recordings in mouse scaffolds under MASH or CTRL conditions. (C) The Amplitude of Ca²⁺ signaling in mice scaffolds induced by ATP was reduced in steatotic scaffolds (p<0.001, n=10-40 cells from three scaffolds in each group). (D) Summary of the ATP-evoked Ca²⁺ profile across the experimental groups. (E) Hematoxylin and eosin-stained sections of control ECM and MASH ECM recellularized with HepG2 cells. Immunohistochemical staining of ITPR1, ITPR2, and ITPR3– positive cells in control ECM and MASH ECM recellularized with HepG2 cells. Scale bars: 50 µm and 20 ∝μ.

In addition to performing calcium assay on rat MASH ECM, we also performed this analysis on human ECM from livers of patients who had MASH and progressed to HCC. Our main objective was to understand whether a diseased human derived ECM could still provide an environment that allows cell development. So, human MASH/neoplastic livers were decellularized (supplementary material) and then, human MASH/neoplastic ECM were also recellularized with HepG2 cells for 14 days and submitted to intracellular Ca²⁺ signaling analysis (Figure 8). Time-lapse confocal images of Fluo-4–loaded HepG2 cells within Human MASH/neoplastic ECM showed that the cells were able to be responsive to ATP stimulation (Figure 8A). We were able to detect specific responses of different cells into the human MASH/neoplastic ECM after ATP stimulation (Figure 8B). We also observed that the profile relates to the Ca²⁺ pattern after ATP stimulation was diverse with distinct cells responding in wave, compound, transient and oscillatory manner (Figure 8C). Immunohistochemistry analysis for intracellular calcium channel isoforms 1, 2 e 3 (ITPR1, 2 e 3) were also evaluated (Figure 8D). All isoforms were identified in HepG2 cultured in human MASH/neoplastic ECM. Together, these results suggest that, although the ECM is diseased, it is still able to promote a physiological environment that ensures cell survival.

**Figure 8.**
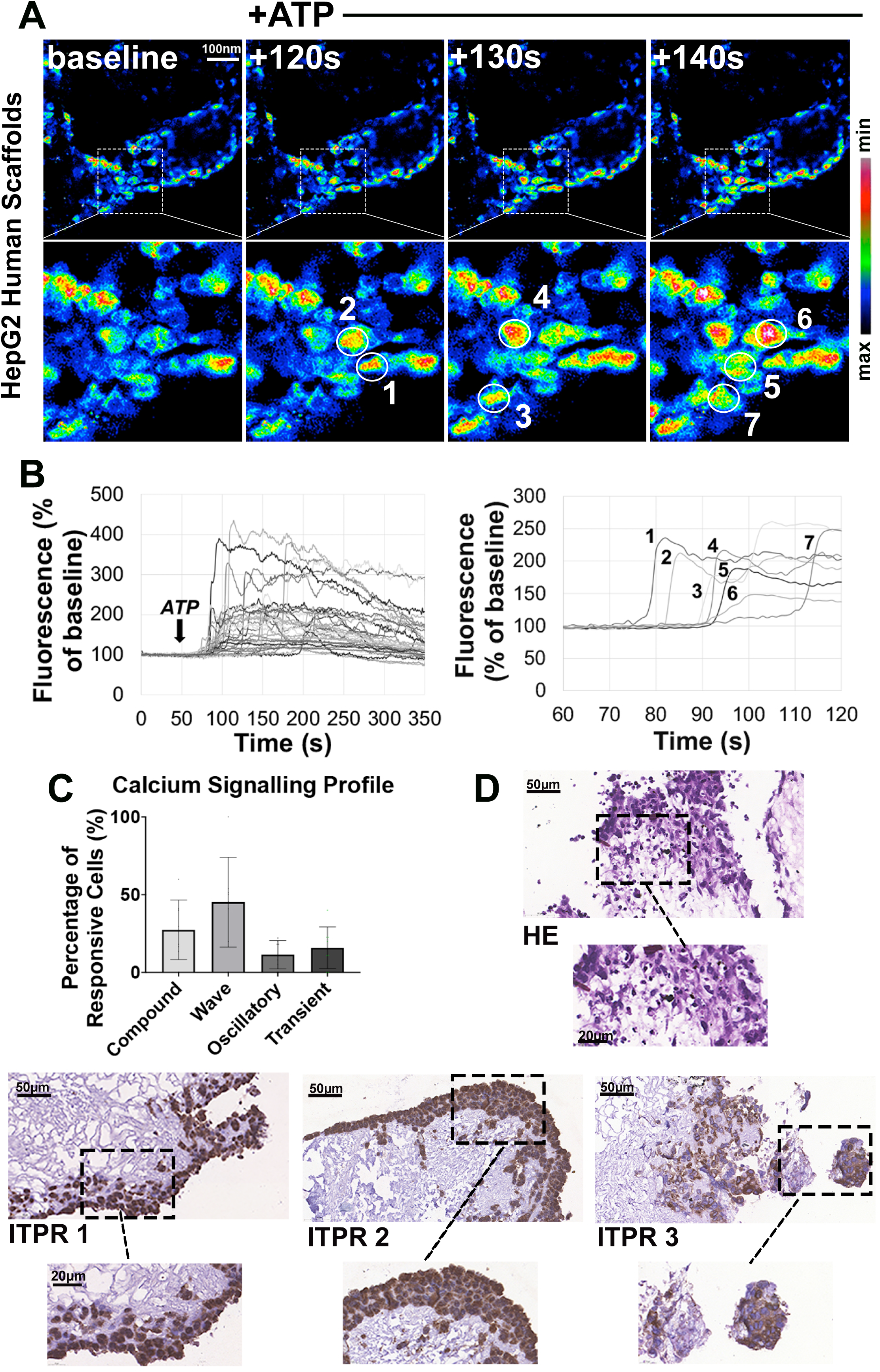
Ca²⁺ dynamics in HepG2-recellularized human scaffolds. (A) Representative time-lapse confocal images of intracellular Ca²⁺ changes (Fluo-4/AM) in HepG2 cells seeded on human decellularized scaffolds. Fluorescence frames are shown at baseline and +120, +130, and +140 s after ATP stimulation. The pseudocolor scale (right) represents the minimum to maximum Ca²⁺ fluorescence intensity. (B) Quantification of ATP-evoked Ca²⁺ signals in HepG2-recellularized human scaffolds, with individual traces representing single-cell recordings. Rescaled representative tracing of individual cells (1–7) that responded sequentially to ATP. (C) Summary of the ATP-evoked Ca²⁺ signaling profile of HepG2 cells seeded on human decellularized scaffolds. (D) Hematoxylin and eosin-stained sections from human MASH/HCC ECM recellularized with HepG2 cells. Immunohistochemical staining of ITPR1, ITPR2, and ITPR3– positive cells in human MASH/HCC ECM recellularized with HepG2 cells. Scale bars: 50 and 20 μm.

## Discussion

In tissue engineering, the emerging concept of “extracellular matrix (ECM) memory” suggests that the ECM preserves information from its tissue of origin and actively influences microenvironmental remodeling based on prior tissue characteristics. The formerly prevailing notion of the ECM as a passive and static scaffold limited to supporting cell adhesion and survival has been fundamentally revised by accumulating evidence demonstrating its active and bidirectional communication with cells in regulating tissue homeostasis [25]. Through direct interactions with cell-surface receptors, the ECM exerts a profound influence on cellular phenotypes. Integrins play a central role in mediating ECM-cell crosstalk by transducing biochemical and biomechanical cues into intracellular signaling, thereby enabling hepatocytes and non-parenchymal cells to dynamically respond to microenvironmental perturbations. Likewise, CD44 interacts with ECM glycosaminoglycans (GAGs), particularly with hyaluronan, and activates signaling pathways involved in inflammatory signaling, tissue repair, and metabolic adaptation. In this setting, constitutional alterations of the ECM, characteristic of all stage liver injuries, can modify ligand-receptor affinity and reshape signal transduction and downstream signaling cascades, ultimately driving intracellular reprogramming linked to the acquirement of disease-associated phenotypes [26]. Moreover, ECM remodeling affects cellular function through indirect mechanisms, as matrisome-associated GAGs bind and sequester cytokines, chemokines, and growth factors, functioning as a reservoir of bioactive mediators that can be rapidly mobilized upon proteolytic activation during tissue injury [25]. Taken together, these observations support the hypothesis that both intrinsic ECM components and retained soluble mediators contribute to the establishment of an ECM memory, enabling sustained and dynamic modulation of the hepatic microenvironment. The concept of ECM memory was previously demonstrated by our group (Dias *et al*., 2023) [7], showing that transplantation of a healthy Acellular Liver Scaffold (ALS) into partially hepatectomized cirrhotic rats led to complete graft recellularization by healthier cells. Notably, the transplanted ALS remained unaffected by the diseased hepatic environment, and although repopulated by cells of cirrhotic origin, it enabled the formation of physiologically normal tissue, providing an effective template for cell growth and supporting the development of functional tissue with normal biochemical, histological, and ultrasonographic features. Based on these findings, the present study sought to expand the understanding of ECM memory through the use of MASH derived ECM.

The rising prevalence of MASH has profoundly affected liver transplantation by reducing the quality of available donor organs and exacerbating organ shortage [26]. Accordingly, in the present study, we chose to employ steatohepatitis models to enhance clinical relevance of our findings. Previously, we demonstrated that transplantation of a healthy ECM can promote a functional healthy tissue recellularization and improve recipient diseased liver function [7]. In the present study, we aimed to advance the translational applicability of this approach by investigating the mechanisms underlying *in vivo* recellularization of diseased ECM within steatotic recipients and assessing the effects of an unhealthy graft on recipient liver physiology. To our knowledge, this is the first study to evaluate a MASH ECM partial transplantation, offering new insights into how scaffold condition affects the liver environment and tissue regeneration. Steatohepatitis induction was performed through a combination of periodic CCl4 injections and a high-fat diet containing palm oil derived fat, fructose, and cholesterol, in accordance with Radhakrishnan *et al*, 2021 considerations regarding disease induction diets on preclinical MASH models [27]. Following disease establishment, confirmed by increased body weight, biochemical parameters evaluation, histology, lipid quantitative analysis and metabolomics, MASH livers were decellularized using a modified detergent-based perfusion protocol adapted from our previously described method, already used in healthy ECM decellularization [7]. To this end, the SDS perfusion duration was extended to overcome the increased decellularization resistance associated with the steatotic ECM, enabling the diseased liver decellularization and successful generation of a MASH ECM.

We observed that the MASH ECM were macroscopically distinct from controls, exhibiting significant heavier weight, histological differences, specially in collagen deposition patterns and increased collagen content along with disorganized three-dimensional fiber structure upon SHG analysis. Those findings were consistent with those reported by Fan *et al* [28], in which the generation of a liver ECM derived from metabolic disease-affected mice revealed a reduced collagen network connectivity and shorter bundles. Biochemically, triglyceride (TGC) assay showed that TGC levels in MASH ECM were comparable to those identified in MASH livers prior to decellularization. This observation is in accordance with findings reported by Acun *et al*, 2024 [15], who demonstrated that TGC content in human steatotic livers does not change significantly following decellularization, supporting the hypothesis that TGC accumulation in steatosis is predominantly retained within the ECM rather than the cellular compartment. The architectural and biochemical remodeling observed in MASH ECM may have direct functional consequences for cells that encounter this diseased microenvironment, since the modified ECM can convey pathological biochemical and biomechanical cues that, when sensed by repopulating cells, may promote cell activation and differentiation. Given this rationale, after *in vivo* recellularization, the identification of positive markers for hepatic stellate cells and myofibroblasts, the two principal effector cell populations responsible for fibrosis generation and maintenance, and the upregulation of *Col1a1,* a key fibrinogenesis regulator, within the recellularized MASH scaffolds suggest that cells shift toward an activated pro-fibrotic phenotype when influenced by MASH ECM. Thus, when cells migrate into a MASH ECM, they are likely to respond to these pathological cues by enhancing ECM synthesis and remodeling, leading to the establishment of a feed-forward loop in which the diseased matrix not only reflects fibrosis but also actively sustains and amplifies it. To further investigate the hypothesis, we next transplanted the MASH ECM into healthy recipients. Remarkably, following recellularization, we observed a similar spatial colocalization of steatosis and collagen fibers, together with confirmed expression of desmin and α-smooth muscle actin, indicating the presence of fibrogenic hepatic stellate cells and myofibroblasts. This encountered pattern closely resembled that observed when MASH ECM was transplanted into MASH recipients. Therefore, the recellularization of MASH ECM was sufficient to induce a steatotic-like tissue phenotype even when transplanted into a healthy recipient. This finding underscores the pivotal role of the diseased ECM in shaping tissue physiology and supports the concept that MASH ECM is not merely a consequence of disease progression, but a fundamental driver of cellular behavior and fibrosis persistence after transplantation.

Following the observation that MASH ECM transplantation in both MASH and healthy recipients results in steatohepatitis maintenance, we next sought to investigate the metabolic profile of the recellularized tissues. Interestingly, this analysis revealed that both MASH and healthy animals transplanted with MASH ECM exhibited similar metabolic signatures. PCA and Venn diagram analyses showed that, in both conditions, MASH-derived ECM induces similar metabolic profiles after transplantation, suggesting that the recipient hepatic microenvironment influences the metabolism of cells within the transplanted ECM, and that the transplanted MASH ECM also directs cellular metabolism, inducing steatohepatitis-like features in resident cells. These profiles were characterized by the upregulation of lipid metabolism related metabolites, including 1-O-hexadecyl-sn-glycero-3-phosphocholine (lyso-PAF) and 4β-hydroxycholesterol 4-acetate (4β-HC-Ac), and glycerophospholipids. Lyso-PAF is a metabolic precursor of the platelet activating factor (PAF), generated when PAF is de-acetylated by PAF acetylhydrolases [30]. Conversely, lyso-PAF can also be re-acetylated back to active PAF by lyso-PAF acetyltransferases, serving as both a stable reservoir of potential PAF and an intermediate in the balance between its synthesis and breakdown [30, 31]. PAF contributes to the development and progression of steatohepatitis by promoting hepatic lipid accumulation and amplifying inflammatory signaling [32]. Activation of the PAF receptor triggers intracellular calcium mobilization, oxidative stress, and insulin resistance, which together disrupt hepatocellular lipid homeostasis and favor triglyceride synthesis and storage [32]. Moreover, PAF enhances immune cell recruitment and cytokine production, thus sustaining chronic inflammation and reinforcing the link between inflammatory signaling and metabolic dysfunction [32]. 4β-HC-Ac is derived from the esterification of 4β-hydroxycholesterol (4β-HC), an endogenous oxysterol that promotes hepatic lipogenesis by acting as a ligand for the liver X receptor (LXR), thereby selectively upregulating the lipogenic transcription factor SREBP1c [32, 33]. Activation of the LXR-SREBP1c axis by 4β-HC enhances the expression of lipogenic genes, stimulating *de novo* triglyceride synthesis and lipid droplet accumulation in hepatocytes, which are central features in the development and progression of steatohepatitis [32]. Finally, glycerophospholipids upregulation suggests an enhanced phospholipase activity that impairs membrane integrity, promotes endoplasmic reticulum and mitochondrial stress, and generates pro-inflammatory lipid mediators [33]. In addition to metabolic profile, MASH ECM also modulates important metabolic pathways such as lipid metabolism, ECM-specific enrichment of arginine and proline metabolism, and arachidonic acid metabolism, pathways closely related to extracellular matrix remodeling, inflammatory signaling, and fibrogenesis. These observations confirmed our molecular and histological findings.

Together, these findings corroborate the hypothesis that MASH ECM functions as a reservoir of steatohepatitis-associated factors capable of inducing a MASH-like metabolic phenotype even when transplanted in a healthy tissue. This concept is further strengthened by the marked metabolic differences observed when healthy animals transplanted with MASH ECM are compared with control animals transplanted with healthy ECM. Therefore, these observations support the notion that MASH ECM retain disease-associated properties and actively modulate their surrounding microenvironment, reinforcing the concept of ECM memory.

After in vivo analysis, we next investigated whether MASH-derived ECM could influence the phenotype and physiology of human liver cells in vitro. Through IHC assessment, both HepG2 cells recellularized within MASH ECM and those exposed to MASH ECM-supplemented medium displayed lipid accumulation patterns comparable to the oleic acid-treated positive control. Accordingly, cells exposed to low-fat control medium exhibited minimal lipid staining, indicating the absence of pathological cues in a health-promoting microenvironment. These findings suggest that not only the MASH ECM but also molecules secreted by the ECM can promote lipid accumulation in HepG2 cells. Given these results, indicative of MASH ECM-induced dysfunction, we next explored the cellular mechanisms underlying MASH phenotype acquisition by assessing intracellular Ca^2+^ dynamics, since Ca^2+^ is a central regulator of hepatic metabolism and cellular homeostasis. Consistent with the diseased morphological phenotype, MASH ECM directly reprograms intracellular Ca^2+^ signaling by altering both signal amplitude and response dynamics. This shift indicates impaired intercellular communication and aberrant Ca^2+^ handling, favoring non-canonical signaling pathways linked to pathological Ca^2+^ homeostasis. These data highlighting the role of diseased ECM in inducing disease-like cellular features are in accordance with Miyauchi *et al,* 2017 [10] results rationale, in which the culture of hepatocarcinoma cells in fibrotic ECM accelerated tumorigenesis process progression when compared with the culture in a control medium. Similarly, Nouri *et al*, 2025 [34] showed that culturing HUVECs and LX-2 cells in fibrotic ECM leads to increased expression of cancer-associated genes relative to control cultures. While previous studies have examined cell culture within healthy or fibrotic liver ECM, our study is, to the best of our knowledge, the first to investigate human liver cell culture within steatotic ECM, broadening the *in vitro* investigation spectrum of liver ECM impact for beyond exclusive oncology associated-questions by now focusing on metabolic disease associations. The integration of evidence from the literature with the results of our animal and human experiments reinforces the hypothesis that tissue phenotype and function are modulated by microenvironment-driven cues.

Understanding how pathological memory is retained and transmitted by diseased ALS is a critical step toward developing strategies to reprogram these scaffolds into healthy-like matrices. The clinical use of healthy-like ALS in liver disease patients could underlie the improvement of diseased organs functionality, likewise we have proposed with our previous findings [7]. Such approaches could enable the reuse of marginal donor livers through decellularization, followed by ECM treatment and pathological memory modification, generating bioengineered grafts capable of promoting tissue repair in recipient damaged organs. The exciting results of our Ca2+ assay evaluating ATP-induced responses in HepG2 cells recellularized within human MASH/HCC-derived ECM, demonstrate that, despite its pathological origin, the ECM can sustain a physiologically permissive microenvironment that supports cell survivals. This finding represents an initial step toward the translational application of human disease-derived ECM as functional bioengineered platforms. Looking ahead, our goal is to reprogram disease-associated ECM memory into a healthy-like ECM capable of promoting physiological cell development and function. Ultimately, this strategy could expand the therapeutic possibilities for patients with end-stage liver disease, improving organ functionality to extend survival while waiting for liver transplantation, and helping to reduce mortality on liver transplant waiting lists. Moreover, modifying disease-associated ALS memory may allow novel liver tissue engineering strategies, facilitating whole ALS recellularization through IPSc-derived liver cells or spheroids, and the consequent creation of functional bioengineered livers, thereby contributing to long-term solutions for organ shortage.

## Conclusion

In conclusion, we show here that ECM obtained from MASH livers retained a memory based on biochemical, and structural cues that promoted steatohepatitis maintenance. Our findings show that ECM obtained from marginal livers should be treated prior to use in liver transplantation to minimize the adverse effects of residual fat in the scaffold.

## Authors’ contributions

Study concept and design: R.C.S.G, M.L.D; Acquisition of data: G.R.P, L.D.G.B, Y.P, A.C.S.F, M.I.M.A.C.G, J.H.O.B, T.O.S, I.Z.L.F.F, B.F.S, M.F.L, H.F.C, C.B.O.A, L.P.G, E.A.A, C.F.P, M.A.A; Analysis and interpretation of data: G.R.P, M.A.A, M.F.L, A.M.G, R.A.S.S, R.C.S.G, M.L.D; Drafting of the manuscript: G.R.P, L.D.G.B, M.A.A, M.L.D; Critical revision of the manuscript for important intellectual content: All authors. Statistical analysis: G.R.P, M.A.A, M.L.D; Obtained funding: R.C.S.G, M.L.D.

## Acknowledgements

We thank João Vitor Santana for his technical assistance during the macrovesicular and microvesicular steatosis quantification, Dr.ª Raiana A. Q. Barbosa for her assistance with PCR steps, Dr. Vitor Pelegati and Dr. Mariana O. Baratti for obtaining the SHG results. We also thank the Rudolf Barth Electron Microscopy Platform of the Oswaldo Cruz Institute for the use of the scanning electron microscope (JEOL-JSM-6390-LV, Akishima, Tokyo, Japan).

## References

1. Devarbhavi H, Asrani SK, Arab JP, Nartey YA, Pose E, Kamath PS. Global burden of liver disease: 2023 update. J Hepatol. 2023;79(2):516–537. doi:10.1016/j.jhep.2023.03.017

2. Terrault NA, Francoz C, Berenguer M, Charlton M, Heimbach J. Liver Transplantation 2023: Status Report, Current and Future Challenges. Clin Gastroenterol Hepatol. 2023;21(8):2150–2166. doi:10.1016/j.cgh.2023.04.005

3. Zamora-Valdes D, Leal-Leyte P, Kim PT, Testa G. Fighting Mortality in the Waiting List: Liver Transplantation in North America, Europe, and Asia. Ann Hepatol. 2017;16(4):480–486. doi:10.5604/01.3001.0010.0271

4. Starzl TE, Groth CG, Brettschneider L, et al. Orthotopic homotransplantation of the human liver. Ann Surg. 1968;168(3):392–415. doi:10.1097/00000658-196809000-00009

5. Organ Procurement & Transplantation Network (OPTN). U.S. Department of Health & Human Services Health Resources and Services Administration. https://optn.transplant.hrsa.gov/. Accessed January 29, 2026.

6. Mir TA, Alzhrani A, Nakamura M, et al. Whole Liver Derived Acellular Extracellular Matrix for Bioengineering of Liver Constructs: An Updated Review. Bioengineering (Basel). 2023;10(10):1126. Published 2023 Sep 25. doi:10.3390/bioengineering10101126

7. Dias ML, Wajsenzon IJR, Alves GBN, et al. Cirrhotic Liver Sustains In Situ Regeneration of Acellular Liver Scaffolds after Transplantation into G-CSF-Treated Animals. Cells. 2023;12(7):976. Published 2023 Mar 23. doi:10.3390/cells12070976

8. Zhang H, Zhang Y, Ma F, Bie P, Bai L. Orthotopic transplantation of decellularized liver scaffold in mice. Int J Clin Exp Med. 2015 Jan 15;8(1):598–606. PMID: 25785034; PMCID: PMC4358489.

9. Naeem EM, Sajad D, Talaei-Khozani T, et al. Decellularized liver transplant could be recellularized in rat partial hepatectomy model. J Biomed Mater Res A. 2019;107(11):2576–2588. doi:10.1002/jbm.a.36763

10. Miyauchi Y, Yasuchika K, Fukumitsu K, et al. A novel three-dimensional culture system maintaining the physiological extracellular matrix of fibrotic model livers accelerates progression of hepatocellular carcinoma cells. Sci Rep. 2017;7(1):9827. Published 2017 Aug 29. doi:10.1038/s41598-017-09391-y

11. Mazza G, Telese A, Al-Akkad W, et al. Cirrhotic Human Liver Extracellular Matrix 3D Scaffolds Promote Smad-Dependent TGF-β1 Epithelial Mesenchymal Transition. Cells. 2019;9(1):83. Published 2019 Dec 28. doi:10.3390/cells9010083

12. Booth AJ, Hadley R, Cornett AM, et al. Acellular normal and fibrotic human lung matrices as a culture system for in vitro investigation. Am J Respir Crit Care Med. 2012;186(9):866–876. doi:10.1164/rccm.201204-0754OC

13. Ogiso S, Yasuchika K, Fukumitsu K, et al. Efficient recellularisation of decellularised whole-liver grafts using biliary tree and foetal hepatocytes. Sci Rep. 2016;6:35887. Published 2016 Oct 21. doi:10.1038/srep35887

14. Uygun BE, Soto-Gutierrez A, Yagi H, et al. Organ reengineering through development of a transplantable recellularized liver graft using decellularized liver matrix. Nat Med. 2010;16(7):814–820. doi:10.1038/nm.2170

15. Acun A, Fan L, Oganesyan R, et al. Effect of Donor Age and Liver Steatosis on Potential of Decellularized Liver Matrices to be used as a Platform for iPSC-Hepatocyte Culture. Adv Healthc Mater. 2024;13(13):e2302943. doi:10.1002/adhm.202302943

16. Goldaracena N, Cullen JM, Kim DS, Ekser B, Halazun KJ. Expanding the donor pool for liver transplantation with marginal donors. Int J Surg. 2020;82S:30–35. doi:10.1016/j.ijsu.2020.05.024

17. Younossi ZM, Golabi P, Paik JM, Henry A, Van Dongen C, Henry L. The global epidemiology of nonalcoholic fatty liver disease (NAFLD) and nonalcoholic steatohepatitis (NASH): a systematic review. Hepatology. 2023;77(4):1335–1347. doi:10.1097/HEP.0000000000000004

18. Golabi P, Paik JM, AlQahtani S, Younossi Y, Tuncer G, Younossi ZM. Burden of non-alcoholic fatty liver disease in Asia, the Middle East and North Africa: Data from Global Burden of Disease 2009-2019. J Hepatol. 2021;75(4):795–809. doi:10.1016/j.jhep.2021.05.022

19. Younossi ZM, Stepanova M, Ong J, et al. Nonalcoholic Steatohepatitis Is the Most Rapidly Increasing Indication for Liver Transplantation in the United States. Clin Gastroenterol Hepatol. 2021;19(3):580–589.e5. doi:10.1016/j.cgh.2020.05.064

20. Thanapirom K, Caon E, Papatheodoridi M, Frenguelli L, Al-Akkad W, Zhenzhen Z, Vilia MG, Pinzani M, Mazza G, Rombouts K. Optimization and Validation of a Novel Three-Dimensional Co-Culture System in Decellularized Human Liver Scaffold for the Study of Liver Fibrosis and Cancer. Cancers. 2021;13(19):4936. doi:10.3390/cancers13194936

21. Qiu B, Simon MC. BODIPY 493/503 Staining of Neutral Lipid Droplets for Microscopy and Quantification by Flow Cytometry. Bio Protoc. 2016;6(17):e1912. doi:10.21769/BioProtoc.1912

22. Liu Z, Wang P, Liu Z, Wei C, Li Y, Liu L. Evaluation of liver tissue extraction protocol for untargeted metabolomics analysis by ultra-high-performance liquid chromatography/tandem mass spectrometry. J Sep Sci. 2021;44:3450–3461

23. Pang Z, Lu Y, Zhou G, et al. MetaboAnalyst 6.0: towards a unified platform for metabolomics data processing, analysis and interpretation. Nucleic Acids Res. 2024;52(W1):W398–W406. doi:10.1093/nar/gkae253

24. Huling J, Götz A, Grabow N, Illner S. GIFT: An ImageJ macro for automated fiber diameter quantification. PLoS One. 2022;17(10):e0275528. Published 2022 Oct 3. doi:10.1371/journal.pone.0275528

25. Arteel GE. Hepatic Extracellular Matrix and Its Role in the Regulation of Liver Phenotype. Semin Liver Dis. 2024;44(3):343–355. doi:10.1055/a-2404-7973

26. Pérez-Escobar J, Huerta-Álvarez A, Castro-Narro GE, Astudillo-Delgado MI, Carpinteyro-Espin P. Era of metabolic dysfunction-associated steatotic liver disease and impact on the liver donor pool. World J Gastroenterol. 2025;31(37):110164. doi:10.3748/wjg.v31.i37.110164

27. Radhakrishnan S, Yeung SF, Ke JY, Antunes MM, Pellizzon MA. Considerations When Choosing High-Fat, High-Fructose, and High-Cholesterol Diets to Induce Experimental Nonalcoholic Fatty Liver Disease in Laboratory Animal Models. Curr Dev Nutr. 2021;5(12):nzab138. Published 2021 Nov 13. doi:10.1093/cdn/nzab138

28. Fan W, Adebowale K, Váncza L, et al. Matrix viscoelasticity promotes liver cancer progression in the pre-cirrhotic liver. Nature. 2024;626(7999):635–642. doi:10.1038/s41586-023-06991-9

29. Tjoelker LW, Wilder C, Eberhardt C, et al. Anti-inflammatory properties of a platelet-activating factor acetylhydrolase. Nature. 1995;374(6522):549–553. doi:10.1038/374549a0

30. Harayama T, Shindou H, Ogasawara R, Suwabe A, Shimizu T. Identification of a novel noninflammatory biosynthetic pathway of platelet-activating factor. J Biol Chem. 2008;283(17):11097–11106. doi:10.1074/jbc.M708909200

31. Yin H, Shi A, Wu J. Platelet-Activating Factor Promotes the Development of Non-Alcoholic Fatty Liver Disease. Diabetes Metab Syndr Obes. 2022;15:2003–2030. Published 2022 Jul 8. doi:10.2147/DMSO.S367483

32. Moldavski O, Zushin PH, Berdan CA, et al. 4β-Hydroxycholesterol is a prolipogenic factor that promotes SREBP1c expression and activity through the liver X receptor. J Lipid Res. 2021;62:100051. doi:10.1016/j.jlr.2021.100051

33. Yamamuro D, Yamazaki H, Osuga JI, et al. Esterification of 4β-hydroxycholesterol and other oxysterols in human plasma occurs independently of LCAT. J Lipid Res. 2020;61(9):1287–1299. doi:10.1194/jlr.RA119000512

34. Nouri K, Piryaei A, Seydi H, et al. Fibrotic liver extracellular matrix induces cancerous phenotype in biomimetic micro-tissues of hepatocellular carcinoma model. Hepatobiliary Pancreat Dis Int. 2025;24(1):92–103. doi:10.1016/j.hbpd.2024.09.003

