## Supplementary material for "Unravelling the memory of the extracellular matrix using MASH-derived decellularized scaffolds"

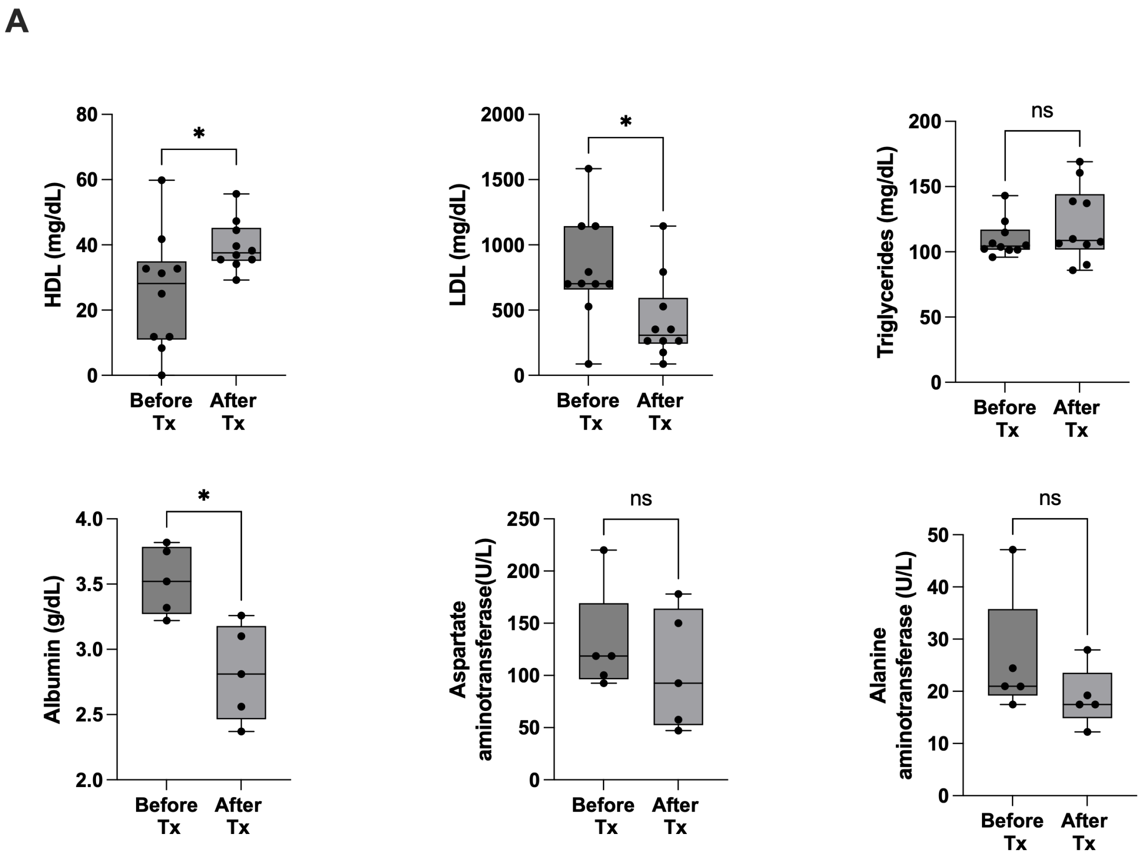


**Supplementary figure 1:** Biochemical analysis of MASH recipients' rats before and 30 days after MASH ECM transplantation. The results are represented as mean ± SD. Statistical significance between the groups was determined using the Student's t-test where *p <0.05, **p <0.01, ***p <0.001, and ****p <0.0001.

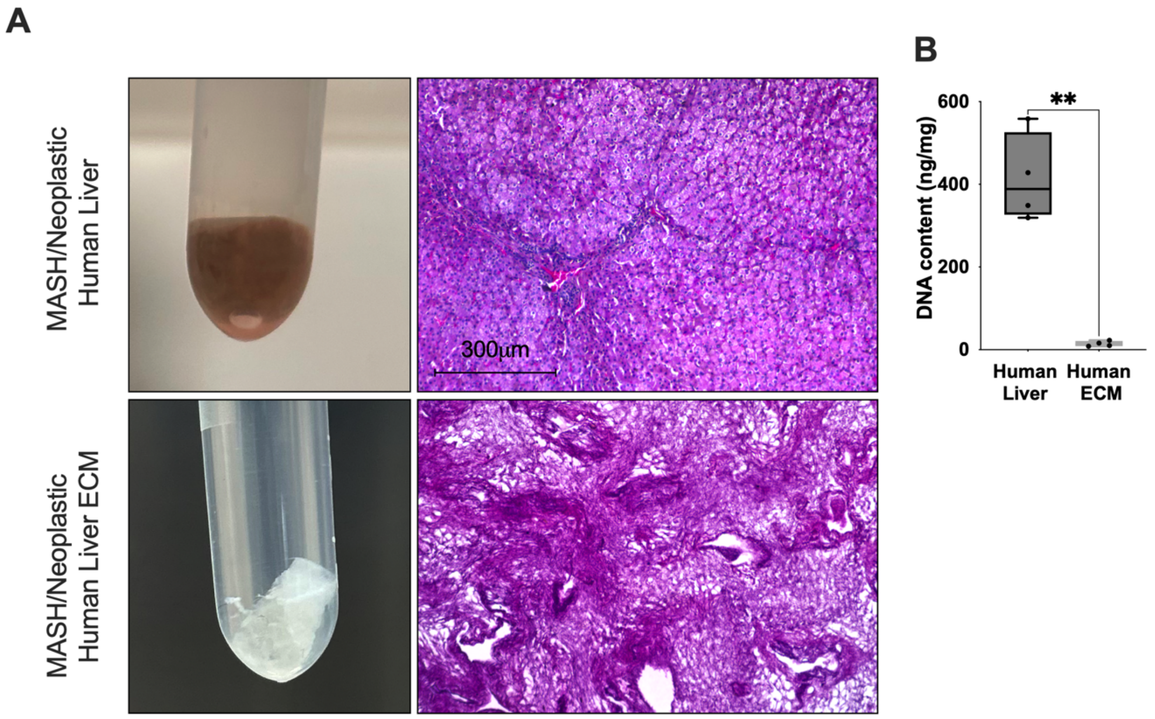


**Supplementary figure 2:** MASH/Neoplastic human liver before and after decellularization. (A) Macroscopic and microscopic views of MASH/Neoplastic human liver before and after decellularization. Histological section stained with H&E obtained from MASH/Neoplastic human liver before and after decellularization. (B) DNA content present in MASH/Neoplastic human liver before and after decellularization. The results are represented as mean ± SD. Statistical significance between the groups was determined using the Student's t-test where $^{**}p < 0.01$.

**Supplementary Table S1:** Forward and reverse primer sequences used for quantitative reverse transcription PCR (qRT-PCR) experiments, including target genes and the endogenous reference gene. All primers were designed to specifically amplify rat transcripts.

| Gene | Forward sequence<br>Reverse sequence | Product size (bp) | Annealing temperature |
|---|---|---|---|
| Acat2 | TCAAGGAATGGGGATGTGGC<br>AAGACAACAAGCAAGCAGGG | 75 | 58ºC |
| Col1a1 | CTCCAGCCGCAAAGAGTC<br>CCCTAAGAGGGAGCAGGAGC | 81 | 58ºC |
| Timp1 | GTCCACAACTGCCAGAACCG<br>AGCAGGGCTCAGATTATGCC | 117 | 60ºC |
| Acaca | GCCTTACAGGATGGTTTGGCCTTT<br>AACAAATTCTGCTGGCGAAGCCAC | 132 | 58ºC |
| PPARα | TTCAATGCCCTCGAACTGGA<br>GCACAATCCCCTCCTGCAAC | 124 | 58ºC |
| Srebp1c | GGCCTGAGAAAGGATGCTCGT<br>CCTCCTGTGTACTTGCCCAT | 95 | 58ºC |
| ApoB | GGACTCATCTGCTACAGCTTAC<br>TGCCCGTGTTCCAATCAA | 95 | 58ºC |
| CPT1α | GAGCCAGACCTTGAAGTACC<br>AGCGACTCTTCAATACTTCCC | 116 | 58ºC |
| GAPDH | TGATTCTACCCACGGCAACT<br>AGCATCACCCCATTTGATGT | 124 | 58ºC |
